# A forward genetic screen identifies Sirtuin1 as a driver of neuroendocrine prostate cancer

**DOI:** 10.1101/2024.08.24.609538

**Authors:** Francisca Nunes de Almeida, Alessandro Vasciaveo, Ainsley Mike Antao, Min Zou, Matteo Di Bernardo, Simone de Brot, Antonio Rodriguez-Calero, Alexander Chui, Alexander L.E. Wang, Nicolas Floc’h, Jaime Y. Kim, Stephanie N. Afari, Timur Mukhammadov, Juan Martín Arriaga, Jinqiu Lu, Michael M. Shen, Mark A. Rubin, Andrea Califano, Cory Abate-Shen

## Abstract

Although localized prostate cancer is relatively indolent, advanced prostate cancer manifests with aggressive and often lethal variants, including neuroendocrine prostate cancer (NEPC). To identify drivers of aggressive prostate cancer, we leveraged *Sleeping Beauty (SB)* transposon mutagenesis in a mouse model based on prostate-specific loss-of-function of *Pten* and *Tp53*. Compared with control mice, *SB* mice developed more aggressive prostate tumors, with increased incidence of metastasis. Notably, a significant percentage of the *SB* prostate tumors display NEPC phenotypes, and the transcriptomic features of these *SB* mouse tumors recapitulated those of human NEPC. We identified common *SB* transposon insertion sites (CIS) and prioritized associated CIS-genes differentially expressed in NEPC versus non-NEPC *SB* tumors. Integrated analysis of CIS-genes encoding for proteins representing upstream, post-translational modulators of master regulators controlling the transcriptional state of *SB*-mouse and human NEPC tumors identified *sirtuin 1* (*Sirt1*) as a candidate mechanistic determinant of NEPC. Gain-of-function studies in human prostate cancer cell lines confirmed that SIRT1 promotes NEPC, while its loss-of-function or pharmacological inhibition abrogates NEPC. This integrative analysis is generalizable and can be used to identify novel cancer drivers for other malignancies.

**Summary:** Using an unbiased forward mutagenesis screen in an autochthonous mouse model, we have investigated mechanistic determinants of aggressive prostate cancer. SIRT1 emerged as a key regulator of neuroendocrine prostate cancer differentiation and a potential target for therapeutic intervention.

## Introduction

The androgen receptor (AR) is the most critical regulator of normal prostate differentiation as well as of all stages of prostate cancer progression (Abate-Shen and Shen, 2000; Gelmann, 2002; Shen and Abate-Shen, 2010). Consequently, prostate cancer treatments have been dominated by approaches to dampen AR signaling (Watson et al., 2015). However, while androgen deprivation therapy (ADT) initially leads to tumor regression, eventually tumors recur as castration-resistant prostate cancer (CRPC), so called because of its continued reliance on AR even in the absence of androgens (Scher and Sawyers, 2005). While further treatment of CRPC with second-generation anti-androgen therapies improves survival, many patients ultimately develop resistance and progress to aggressive disease variants that may no longer be dependent on AR (Watson et al., 2015). It is now understood that the emergence of such aggressive prostate cancer variants, including neuroendocrine prostate cancer (NEPC), occurs via lineage plasticity (Beltran et al., 2016; Ku et al., 2017; Mu et al., 2017; Zou et al., 2017), which refers to the transition from one differentiated cell state to an alternative state (Le Magnen et al., 2018). Therefore, elucidating the mechanisms that preside over the emergence of aggressive prostate cancer should improve treatments by abrogating plasticity-associated drug resistance.

We sought to identify causal drivers of aggressive prostate cancer by implementing a *Sleeping Beauty (*SB) forward genetics mutagenesis screen (Dupuy et al., 2005; Starr et al., 2009). *Sleeping Beauty* is a two-component system consisting of a transposon, which is a DNA element that can randomly move around the genome, and a transposase, which is an enzyme that promotes transposon excision and random insertion elsewhere in the genome. Herein, we refer to the bi-allelic mice having both the transposon and the transposase as the “active SB” mice (*SB+*), and we distinguish these from mono-allelic mice having the transposase but lacking the transposon, which we refer to as the “inactive SB” (*SB–* mice). When both the transposase and transposon are present, as occurs in the *SB+* mice, the transposon can insert randomly into the genome causing mutations; this is not the case in the *SB–* mice, which lack the transposon. When random insertions occur in or near cancer-relevant genes, thereby leading to their aberrant activation or inactivation, such transposon-induced mutations may have cancer-promoting or cancer-modulating effects.

Unlike CRISPR or RNAi based screens, which target known genes and usually in a defined direction *(i.e.*, either gain- or loss-of-function), *SB*-mediated transposition is an unbiased approach in which mutations can occur anywhere in the genome and may result in either gain- or loss-of-function of nearby genes (Beckmann and Largaespada, 2020; Copeland and Jenkins, 2010). Furthermore, since mutagenesis occurs in autochthonous mice, the ensuing mutations arise somatically during the natural time-course of disease progression and in the context of the native tumor environment, analogous to the accumulation of mutations during cancer progression in humans. These features make *SB-*mediated mutational screens attractive for elucidating new mechanisms of cancer progression, as evidenced by several studies (Dupuy et al., 2009), especially in colorectal cancer (Iida et al., 2024; March et al., 2011; Shimomura et al., 2023; Starr et al., 2009; Starr et al., 2011), but also in tumors of the central nervous system (Beckmann et al., 2019), pancreas (Mann et al., 2012), melanoma (Mann et al., 2015), and prostate (Ahmad et al., 2016; Rahrmann et al., 2009). However, *SB-*mediated mutagenesis has been less widely used, compared to RNAi or CRISPR screens, in part due to inherent difficulties in identifying the genes that are responsible for the resulting phenotypes, precisely because of the random nature of the transposon-induced mutagenesis.

In the current study, we have implemented an *SB* screen using a well-characterized genetically engineered mouse model (GEMM) of prostate cancer based on haploinsufficient loss of *Nkx3.1* and homozygous loss-of-function of both *Pten* and *Trp53* (*NPp53* mice (Zou et al., 2017)), which are prevalent genetic alterations in advanced prostate cancer in humans (Abida et al., 2019). Compared with the control *SB*-inactive *NPp53*-*SB(–)* mice, the experimental *SB*-active *NPp53*-*SB(+)* mice develop more aggressive prostate cancer phenotypes with a high incidence of NEPC. Using an integrative systems biology approach—combining phenotypic, transcriptomic, and genomic analyses with state-of-the-art network-based algorithms—we have identified and validated mechanistic determinants of NEPC. Among these was the nicotinamide adenosine dinucleotide (NAD)-dependent deacetylase *sirtuin 1* (*Sirt1*). We show that expression of Sirt1 promotes NEPC while its depletion or pharmacological inhibition dampens trans-differentiation to an NEPC phenotype. Overall, our findings introduce a highly generalizable computational and experimental approach for integrating *SB* mutagenesis with phenotypic and transcriptomic profiles to identify novel cancer drivers.

## Results

### Overall strategy

The strategy to identify mechanistic determinants of advanced prostate cancer utilizing a *Sleeping Beauty (SB)* mutagenesis screen entails integration of phenotypic, transcriptomic, and genomic analyses using state-of-the-art systems biology algorithms with cross-species comparative analyses of mouse and human tumors to identify and prioritize candidate genes, which are then functionally validated (Fig. 1).

**Figure 1.**
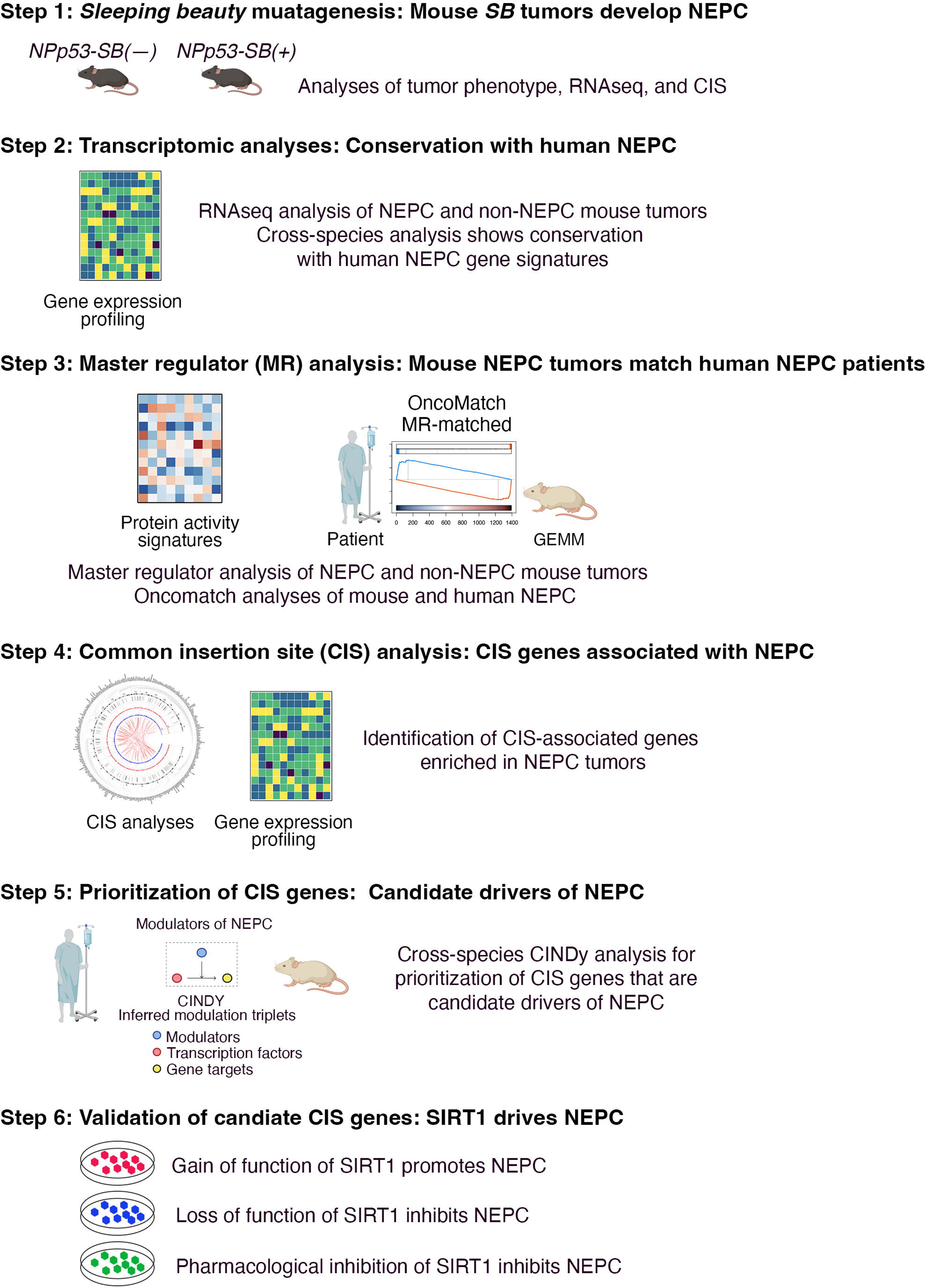
Strategy to identify drivers of aggressive prostate cancer from a *Sleeping Beauty* mutagenesis screen. Shown are the steps undertaken to identify regulators of neuroendocrine prostate cancer (NEPC) based on analyses of *Sleeping Beauty* mutagenesis screen in a genetically engineered mouse model of prostate cancer. As detailed in the text, *SB* mice were generated, and phenotypic and molecular analyses were performed on prostate tumors from these mice, and in comparison with human NEPC. Common insertion sites (CIS) for the transposons were determined, and CIS-associated genes were identified and prioritized based on their expression in NEPC and using the CINDy algorithm to predict mechanistic determinants. Candidate CIS-genes were validated in human prostate cancer models.

*First*, to implement the *SB* screen, we generated the relevant mouse alleles by crossing bi-allelic *SB* mice (Dupuy et al., 2005; Starr et al., 2009) with the *NPp53* mice (Zou et al., 2017) to generate the experimental *NPp53-SB(+)* and control *NPp53-SB(–)* mice (Fig. 1, Step 1). Phenotypic analyses revealed that, compared to control mice, the *NPp53-SB(+)* mice had more aggressive prostate cancer phenotypes, with greater occurrence of metastasis, and a high incidence of NEPC.

*Second*, we performed transcriptomic analyses of the histologically characterized NEPC and non-NEPC prostate tumors from the *SB* mice. This confirmed the expression of established NEPC markers and revealed transcriptional signatures that significantly recapitulated those of human NEPC (Fig. 1, Step 2).

*Third*, we used the VIPER algorithm, which effectively transforms RNA-seq profiles into protein activity profiles (Alvarez et al., 2016), to analyze the transcriptomic data from *SB* tumors with the objective of identifying master regulator (MR) proteins that control the state of NEPC vs. non-NEPC mouse tumors (Fig. 1, Step 3). We then used the extensively validated OncoMatch algorithm (Alvarez et al., 2018; Mundi et al., 2023; Vasciaveo et al., 2023) to assess the fidelity of these murine models to human NEPC and non-NEPC tumors—from relevant publicly available human prostate cancer cohorts— based on MR signature overlap. This revealed that the mouse NEPC tumors represent highly concordant (i.e., high-fidelity) models of human NEPC patient tumors.

*Fourth*, we performed genomic sequencing of NEPC and non-NEPC prostate tumors from the *SB* mice to identify transposon integration sites (Fig. 1, Step 4). TAPDANCE analysis (Sarver et al., 2012) identified common insertion sites (CIS) as well as corresponding CIS-associated genes. Since we expect that the effect of functionally relevant transposons is largely mediated by changes in the expression of the associated gene, we prioritized CIS-associated genes based on their differential expression in NEPC versus non-NEPC tumors. For simplicity, we will use the term CIS-genes to indicate CIS-associated genes that are differentially expressed in NEPC vs. non-NEPC tumors.

*Fifth*, we further prioritized CIS-genes as candidate mechanistic determinants of NEPC transcriptional state using a cross-species approach (Fig. 1, Step 5). Specifically, as proposed in (Chen et al., 2014; Paull et al., 2021), we reasoned that mutations in genes encoding for proteins representing upstream modulators of cell-state relevant MRs may identify mechanistic determinants of NEPC. We thus leveraged the CINDy algorithm (Giorgi et al., 2014; Paull et al., 2021) to systematically identify candidate upstream modulators of NEPC MR activity and prioritized those harboring a CIS event based on the integrated analysis statistical significance (see methods).

*Lastly*, we validated the most likely candidate, *Sirt1*, in human prostate cancer cell models using gain- and loss-of-function studies, as well as pharmacological inhibition to demonstrate its relevance for promoting NEPC (Fig. 1, Step 6). Each of these steps is described in detail below.

### *Sleeping Beauty* mutagenesis accelerate prostate cancer progression and promotes NEPC

We implemented the *SB* screen (Fig. 1, Step 1) using the *NPp53* mice (for *<u>N</u>kx3.1^CreERT2/+^; <u>P</u>ten^flox/flox^; Tr<u>p53</u>^flox/flox^*) (Fig. 2A), a well-characterized genetically engineered mouse model (GEMM) based on combined loss-of-function of *Pten* and *Trp53* specifically in the prostate epithelium (Zou et al., 2017). *NPp53* mice display poorly differentiated prostate cancer that progresses to CRPC; further treatment of these mice with second generation anti-androgens results in progression to more aggressive prostate cancer phenotypes, including NEPC (Zou et al., 2017). Considering their capability to develop aggressive prostate cancer phenotypes, we reasoned that the *NPp53* mice would serve as an excellent model to identify drivers of aggressive prostate cancer, including NEPC. As a control, we used the *NP* mice (for *<u>N</u>kx3.1^CreERT2/+^; <u>P</u>ten^flox/flox^*; (Floc’h et al., 2012)), which develop well-differentiated prostate adenocarcinoma that develops CRPC following androgen deprivation but does not progress to more advanced phenotypes following treatment with second generation anti-androgens (Zou et al., 2017). Tumor induction in the prostatic epithelium of the *NPp53* and *NP* mice is spatially and temporally controlled by activation of the *Nkx3.1^CreERT2^* allele using tamoxifen (Wang et al., 2009b), which also renders haploinsufficient loss of *Nkx3.1,* as is prevalent in human prostate cancer (Cancer Genome Atlas Research, 2015).

**Figure 2.**
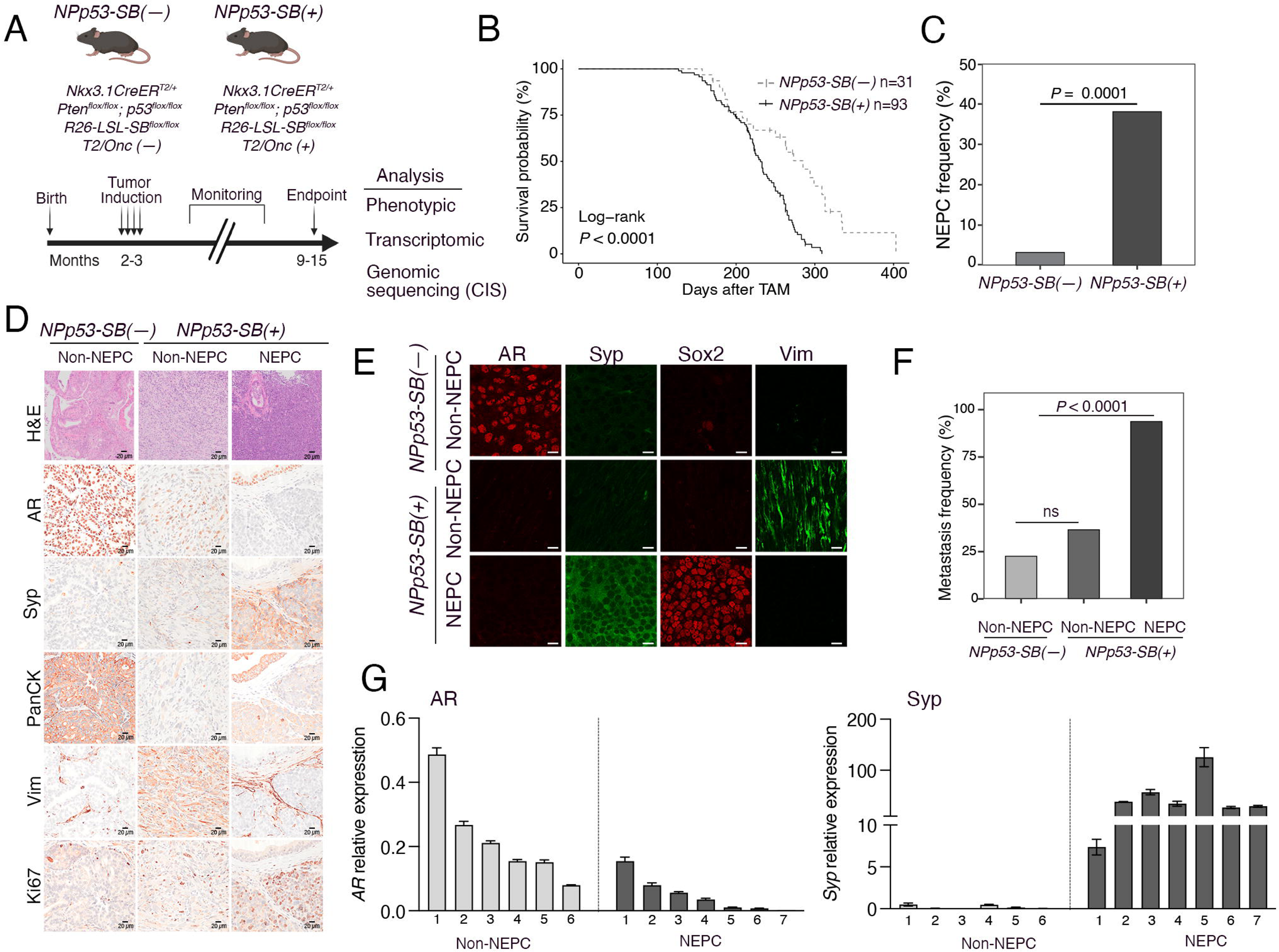
*Sleeping Beauty* mutagenesis of *NPp53* mice promotes neuroendocrine prostate cancer (NEPC). A. <u>Strategy</u>: *NPp53-SB(–)* and *NPp53-SB(+)* mice were induced to form tumors at 2-3 months of age and monitored for up to 15 months, following which tissues were collected for histological, RNA and DNA sequencing analyses. **B.** Kaplan-Meier survival analysis of *NPp53(SB–)* (n=31) and *NPp53(SB+)* (n=93) mice. The *P* value was calculated using a Log-rank test. **C.** NEPC frequency in *NPp53-SB(–)* (n=31) and *NPp53-SB(+)* (n=84) prostate tumors. The *P* value was calculated using the Fisher’s Exact Test. **D,E.** <u>Histopathological analyses.</u> Representative images of prostate tumors from the *NPp53-SB(–)* and *NPp53-SB(+)* mice, showing examples of neuroendocrine prostate cancer (NEPC) and non-NEPC (adenocarcinoma and sarcomatoid) phenotypes. Shown are hematoxylin and eosin (H&E) staining, and immunohistochemical (IHC) or immunofluorescence (IF) staining for markers of prostate cancer and NEPC, namely, AR, Synaptophysin (Syp), Pan-cytokeratin (Pan-CK), Vimentin (Vim), Ki67, and Sox2. Scale bars represent, 20 µm for H&E, 10 µm for IHC and IF images. F. Metastasis frequency in *NPp53-SB(–)* (n=31) and *NPp53-SB(+)* (n=84) mice comparing non-NEPC (n=52) and NEPC (n=32) tumors. **G.** Quantitative real-time PCR showing expression levels of *AR* and *synaptophysin (Syp)* in non-NEPC (n=6) and NEPC (n=7) *NPp53-SB(+)* tumors. See also Figures S1A, S1B, Table 1, and Table S1.

**Table 1:** Summary of the mouse strains used in this study and their associated phenotypes.

| Name | Genotype | N | Phenotype | Metastasis | NEPC |
| --- | --- | --- | --- | --- | --- |
| NPp53-SB(-) | *Nkx3.1*<sup>CreERT2/+</sup>; *Pten*<sup>flox/flox</sup>;<br>*Trp53*<sup>flox/flox</sup>; R26-LSL-<br>*SB11*<sup>flox/flox</sup> | 31 | Poorly differentiated<br>adenocarcinoma with sarcomatoid<br>features | 7/31<br>(22%) | 1/31<br>(3%) |
| NPp53-SB(+) | *Nkx3.1*<sup>CreERT2/+</sup>; *Pten*<sup>flox/flox</sup>;<br>*Trp53*<sup>flox/flox</sup>; R26-LSL-<br>*SB11*<sup>flox/flox</sup>; T2/*Onc2* | 93 | Poorly differentiated<br>adenocarcinoma with sarcomatoid<br>features and neuroendocrine<br>differentiation | 57/93<br>(61%) | 32/84<br>(38%) |
| NP-SB(-) | *Nkx3.1*<sup>CreERT2/+</sup>; *Pten*<sup>flox/flox</sup>;<br>R26-LSL-*SB11*<sup>flox/flox</sup> | 12 | High-grade PIN, well-differentiated<br>adenocarcinoma | 0/12<br>(0%) | 0/12<br>(0%) |
| NP-SB(+) | *Nkx3.1*<sup>CreERT2/+</sup>; *Pten*<sup>flox/flox</sup>;<br>R26-LSL-*SB11*<sup>flox/flox</sup>;<br>T2/*Onc2* | 18 | High-grade PIN, well-differentiated<br>adenocarcinoma | 0/18<br>(0%) | 0/18<br>(0%) |

The *Sleeping Beauty (SB)* system is comprised of two alleles: a conditionally activatable transposase located in the *Rosa26* locus (*Rosa26^LSL-SB11^*) and a transgenic allele expressing the transposon (*T2/Onc2*) (Dupuy et al., 2005; Starr et al., 2009). Cre activation (in this case via the *Nkx3.1^CreERT2^* allele) leads to excision of the *Stop* cassette that blocks expression of the *SB11* transposase, thereby resulting in expression of the transposase, which allows it to bind to terminal repeats at both ends of the *T2/Onc2* transposon to promote its excision and random integration in the genome (Dupuy et al., 2005; Starr et al., 2009). To generate the *SB* mice, we crossed the *Rosa26^LSL-SB11^* and*T2/Onc2* alleles with the *NPp53* mice to generate the experimental *NPp53-SB(+)* mice (for *Nkx3.1^CreERT2/+^; Pten^flox/flox^; Trp53^flox/flox^; Rosa26^LSL-SB11/LSL-SB11^; T2Onc2^Tg/+^)*, which have both the transposase and the transposon, and the control *NPp53-SB(–)* mice (for *Nkx3.1^CreERT2/+^; Pten^flox/flox^; Trp53^flox/flox^; Rosa26^LSL-SB11/LSL-SB11^*), which lack the transposon (Fig. 2A). Tumor induction of the *NPp53-SB(+)* mice results in activation of the transposase and consequent transposition of *T2/Onc2*, as confirmed by PCR analyses (Fig. S1A). In contrast, the *NPp53-SB(–)* mice lack the transposon, and therefore have no transposition events (Fig. S1A). We also generated the corresponding *NP-SB(–)* mice (*for Nkx3.1^CreERT2/+^; Pten^flox/flox^; Rosa26^LSL-SB11/LSL-SB11^*) and the *NP-SB(+)* mice (*for Nkx3.1^CreERT2/+^; Pten^flox/flox^; Rosa26^LSL-SB11/LSL-SB11^; T2Onc2^Tg/+^)* (Fig. S1B,C).

We analyzed mice that had undergone tamoxifen-mediated tumor induction at 2-3 months of age and were monitored for up to 12 months or until they succumbed to prostate cancer, after which their prostate tumors and other tissues were collected for phenotypic and molecular analyses (Fig. 2A). Compared with the *NPp53-SB(–)* mice, the *NPp53-SB(+)* mice had a greater tendency to develop more aggressive prostate cancer phenotypes as evident by their histological phenotype, reduced survival, and increased incidence of metastasis (Fig. 2B-F, Figs S1D-H, Fig. S2A-F, Table 1, Table S1). In particular, the *NPp53-SB(+)* mice displayed a significant reduction in survival relative to the *NPp53-SB(–)* mice (log rank p-value < 0.001; Fig. 2B). In contrast, the *NP-SB(+)* mice did not display a significant decrease in survival or an acceleration of their prostate cancer phenotype, compared with the *NP-SB(–)* mice (Fig. S1B,C; Table S1), consistent with previous observations that they lack the ability to progress to advanced prostate cancer (Zou et al., 2017).

As is characteristic of the *NPp53* mice (Zou et al., 2017), histopathological analyses of prostate tumors from the *NPp53-SB(–)* and *NPp53-SB(+)* mice revealed a range of phenotypes with significant heterogeneity (Fig. 2D, Fig. S2A-F, Table 1, Table S1). However, whereas the spectrum of histopathology of the *NPp53-SB(–)* tumors consisted predominantly of squamous or sarcomatoid phenotypes, a significant percentage of *NPp53-SB(+)* tumors displayed histopathological features of NEPC, which were rarely seen in the *NPp53-SB(–)* tumors (Fig. 2D,E, Fig. S2A-F, Table 1, Table S1). These features include: *(i)* a nesting trabecular growth pattern, *(ii)* small tumor cells with scant cytoplasm, and *(iii)* high mitotic count (Ittmann et al., 2013; Shappell et al., 2004).

The NEPC phenotypes of these *NPp53-SB(+)* tumors were further evident by expression of known markers of NEPC including synaptophysin (Syp), chromogranin A (ChgA), and Sox2, and a corresponding decrease in expression of AR (Fig. 2D,E,G, Fig. S1F). In addition, although their tumor weights were comparable (Fig. S1D), NEPC tumors from the *NPp53-SB(+)* mice were significantly more proliferative compared with non-NEPC tumors from the *NPp53-SB(+)* mice or with tumors from the *NPp53-SB(–)* mice, as evident by quantification of Ki67 immunostaining (Fig. 2D, S1E). Notably, *NPp53-SB(+)* mice displayed a significant increase in the incidence of metastasis, including to the lymph nodes, lung and liver, nearly all of which occurred in the *NPp53-SB(+)* mice with NEPC tumors (Fig. 2F, Fig. S1G,H; S2B; Table 1). Therefore, *SB* mutagenesis of *NPp53* tumors promotes highly aggressive and metastatic prostate cancer with a high incidence of NEPC. Notably, the *NPp53-SB(+)* mice had not undergone hormone deprivation or any other treatment; therefore, *SB* mutagenesis accelerates the development of NEPC that arises *de novo* in hormonally intact untreated mice.

### NEPC tumors from *NPp53-SB(+)* mice are well-conserved with human NEPC

To characterize their transcriptomic phenotype (Fig. 1, Step 2), we performed RNA-sequencing of *SB* tumors that were histologically identified as NEPC or non-NEPC from the *NPp53-SB(+)* and *NPp53-SB(–)* mice (n = 27 and n = 8, respectively; Fig. 3A-C; Table S1, S2). As evident from both differential gene expression analyses and principal-component analysis (PCA), the NEPC *NPp53-SB(+)* tumors clustered separately from both the non-NEPC *NPp53-SB(+)* tumors and the *NPp53-SB(–)* tumors (Fig. 3A,B), indicating that they are molecularly distinct.

**Figure 3.**
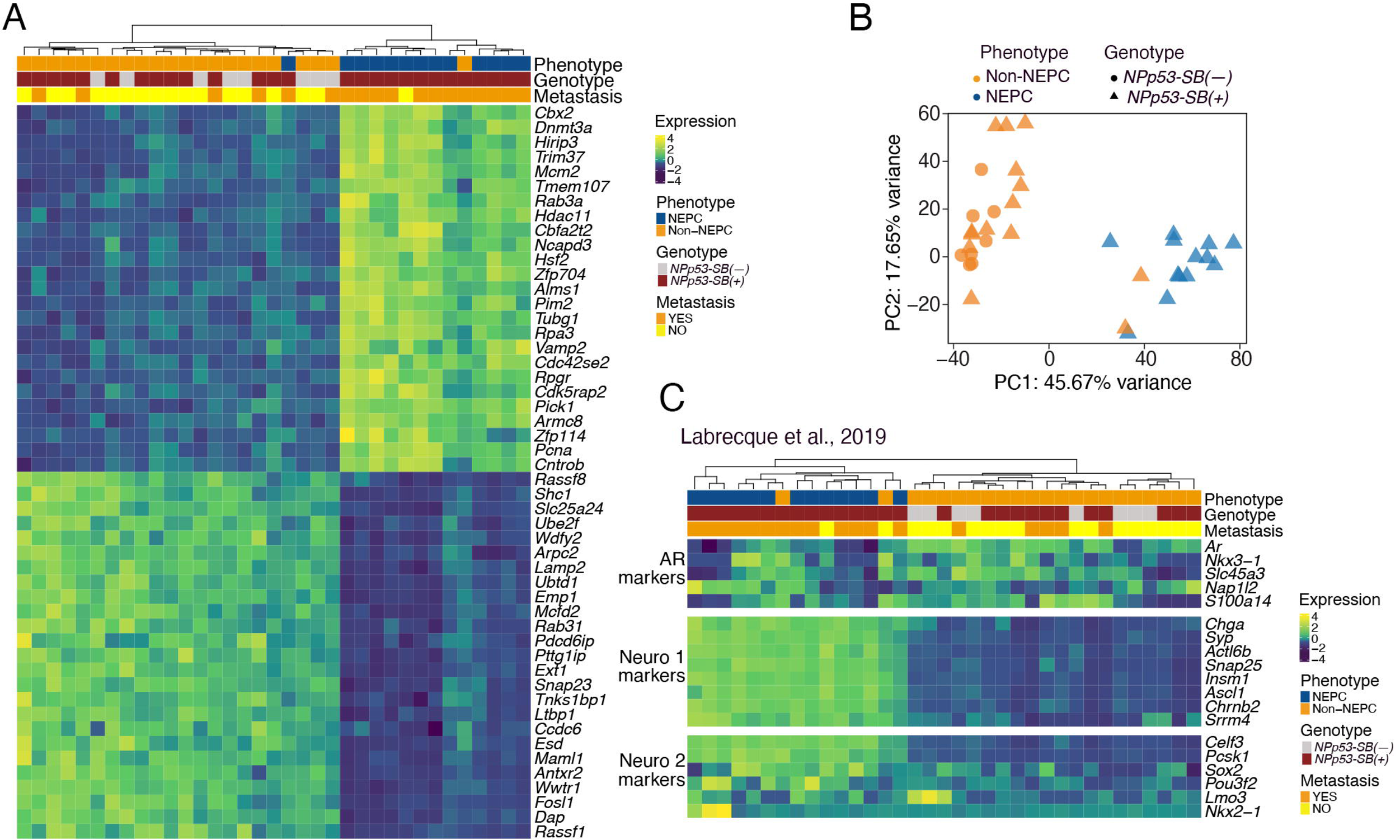
Transcriptomic analysis of *Sleeping Beauty* mouse prostate tumors. A. Unsupervised clustering of RNA seq data visualized by heatmap analyses. Shown are the top 25 and bottom 25 differentially expressed genes between NEPC and non-NEPC tumors. **B.** Principal component analyses of non-NEPC (n=22) and NEPC (n=13) tumors from *NPp53-Ctrl* (n=8) and *NPp53-SB* (n=27) mice. **C.** RNA-seq heatmap showing relative expression levels for genes related to AR and Neuroendocrine markers from the datasets published in (Labrecque et al., 2019). Relevant variables such as phenotype, genotype and metastatic status are shown at the top. See also Table S2.

To determine whether the NEPC tumors from *NPp53-SB(+)* mice recapitulated key molecular features of human NEPC, we assessed the enrichment of gene signatures associated with well-characterized androgen receptor-dependent (AR) or neuroendocrine (NE) sub-types of CRPC (Labrecque et al., 2019) in genes that were differentially expressed in the *SB* mouse tumors (Fig. 3C). These previously characterized signatures comprise a set of AR-regulated genes, as well as NE-associated genes in CRPC subtypes, including a NEURO I panel, comprising REST-repressed genes, and a NEURO II panel comprising transcription factors that regulate NE differentiation (Labrecque et al., 2019). Unsupervised cluster analysis showed that the *SB* tumors that were histologically identified as NEPC overexpressed NEPC-related NEURO I and NEURO II gene sets while having reduced expression of AR-related genes (Fig. 3C). The opposite was observed for the non-NEPC *SB* tumors (Fig. 3C).

We next sought to identify Master Regulator (MR proteins)—representing key drivers of the NEPC phenotype—by analyzing *NPp53-SB(+)* transcriptomes with the VIPER algorithm (Fig. 1, Step 3). For this purpose, we leveraged a previously generated mouse prostate cancer interactome (Vasciaveo et al., 2023), reverse engineered by analyzing large-scale, tissue-specific RNA-seq profiles using the ARACNe algorithm (Basso et al., 2005). These analyses transformed the differential expression of NEPC versus non-NEPC *SB* tumors to VIPER-assessed differential protein activity profiles (Table S3). Protein activity-based cluster analysis identified 3 clusters, corresponding to 3 molecularly distinct tumor subtypes (*C_1_* – *C_3_*) comprising either NEPC or non-NEPC tumors (Fig. 4B, Table S4). Specifically, subtypes *C*_1_ and *C*_2_ mainly comprised *NPp53-SB(–)* tumors and non-NEPC *NPp53-SB(+)* tumors, while subtype *C_3_* comprised the NEPC *NPp53-SB(+)* tumors.

**Figure 4.**
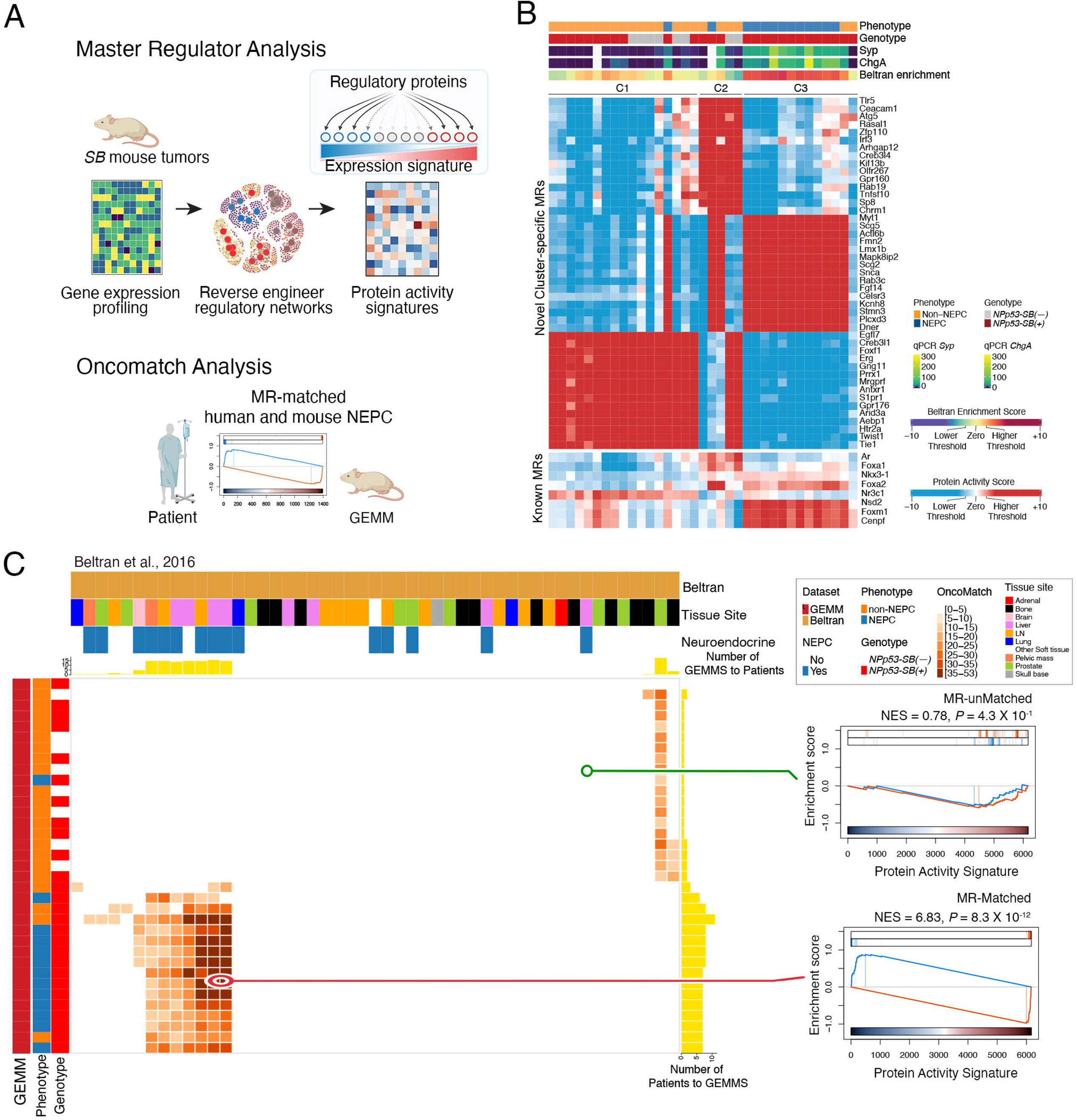
Master regulator profiles of mouse NEPC show conservation with human NEPC tumors. A. <u>Strategy</u>. Master regulators (MRs) of NEPC were determined using VIPER to interrogate mouse prostate tumor regulatory network based on differential gene expression signatures of non-NEPC and NEPC *SB* tumors. To compare with human prostate cancer, the mouse MR signatures were compared with individual patient samples from relevant human cohorts using OncoMatch. **B**. Heatmap of protein activity-based cluster analysis of non-NEPC and NEPC *SB* mouse prostate tumors. Projected are the top-15 MRs that distinguish each cluster. Also shown are the activity levels of known MRs of advanced prostate cancer and NEPC. Shown at the top are the phenotype and genotype, quantitative real-time gene expression levels for *synaptophysin* (*Syp*) and *chromograninA* (*ChgA*), and enrichment of the Beltran NEPC features. **C.** OncoMatch heatmap representing conservation of master regulators (MRs) between individual *SB* tumors (rows, n=32) and patient tumors (columns, n=49) from the Beltran cohort. Patient variables (*i.e.* assessment of neuroendocrine features and tissue site) are shown on the top horizontal bars, while GEMM variables (*i.e.* phenotype and genotype) are shown on the left side vertical bars. Yellow bars at the top of the heatmaps show the number of GEMMs that match each patient, while the yellow bars on the right side show the number of patients that match each GEMM. The GSEA on the left show representative *SB* tumor and human patient pairs showing an example of an MR-unmatched (low-fidelity, top) and an MR-matched (high-fidelity, bottom) pair. See also Fig.S3, Tables S3, S4, and S5.

Notably, tumors in *C_3_* have robust expression of established NEPC markers, including synaptophysin and chromogranin A (Fig. 4B, top legend), as evidenced by qPCR (see Fig. 2G, S1F). Furthermore, a gene panel representing a human NEPC signature—as assessed in a well-characterized human prostate cancer cohort comprised of adenocarcinoma and NEPC from mCRPC patients (*n*=49), herein named Beltran cohort (Beltran et al., 2016)—were highly enriched in genes differentially expressed in *C*_3_ tumors vs. *C*_1_ and *C*_2_ tumors (Fig. 4B, top legend), as determined by protein activity signatures of *C_3_* tumors as assessed by Gene Set Enrichment Analysis (GSEA).

The proteins most differentially active in *C_3_* tumors, representing candidate MRs, include transcription factors that have been previously reported as NEPC related, such as Myt1 (Fig. 4B) (Guo et al., 2019). Interestingly, among the top 15 MRs in *C_3_*, 13 are known to be expressed in the nervous system and/or involved in nervous system development, including the homeobox gene, Lmx1b, the Notch-binding receptor, Dner, the regulator of synaptic transmission or maturation, Snca and Rab3c (e.g., (Ding et al., 2003; Fischer von Mollard et al., 1994; Han et al., 2021; Hsieh et al., 2013; Siddiqui et al., 2016)). In addition to these novel MRs, C_3_ also presented high activity of MRs known to be associated with aggressive prostate cancer, including FoxM1, Cenpf, and Nsd2, as well as MRs specifically associated with NEPC, including Foxa2 and Foxa1 (Fig. 4B) (Han et al., 2022; Vasciaveo et al., 2023).

Next, we determined whether MR signatures of the mouse NEPC tumors are conserved with analogous MRs from human NEPC patients (Fig. 4A,C, Fig. S3, Table S5). For comparative analysis with the mouse signatures (described above), we analyzed two well-characterized human cohorts that include NEPC patient samples. In particular, we queried the previously discussed Beltran cohort (Beltran et al., 2016)—comprising n = 34 CRPC and n = 15 NEPC tumors—as well as a second cohort comprising of post-treatment metastatic biopsies from mCRPC patients collected as part of the Stand Up to Cancer-Prostate Cancer Foundation study (SU2C) (Abida et al., 2019)—comprising 210 non-NEPC, 22 NEPC and 34 unclassified tumors. Similar to the mouse analysis, we used VIPER to transform the transcriptional profile of each patient-derived sample in these cohorts into a protein activity profile (Table S5), based on a previously generated human prostate cancer interactome (Vasciaveo et al., 2023).

We then assessed the enrichment of the 25 most activated (25↑) and 25 most inactivated (25↓) candidate MRs of each human tumor in proteins differentially active in each *SB* tumor (Table S5). High-fidelity (*i.e.,* well-conserved) *SB* tumors were identified using the OncoMatch algorithm as those with highly conservative statistically significant enrichment (*P* ≤ 10^-5^, by Bonferroni corrected 1-tail aREA test) (Vasciaveo et al., 2023). We used a fixed number of MR proteins (i.e., 25↑+25↓) since: (i) this is required to make the statistics of enrichment analyses comparable across proteins and cohorts; and (ii) we have previously shown that an average of 50 MRs is sufficient to account for the canalization of functionally-relevant genetic alterations (Paull et al., 2021; Vasciaveo et al., 2023). These analyses revealed that each individual NEPC mouse tumor was a faithful model for at least one NEPC patient (Fig. 4C, Fig. S3). Analyses of the Beltran cohort revealed that 66.7% of the NEPC patients matched to at least one mouse NEPC tumor (Fig. 4C). Similarly, the SU2C cohort revealed a 45.5% match between NEPC patients and NEPC mouse tumors (Fig. S3). Taken together, these findings show that the NEPC tumors from the *SB* mice are high-fidelity models of human NEPC.

### Identification of CIS-associated genes enriched in NEPC

To identify genes associated with NEPC in the *SB* tumors, we; *(i)* first performed genomic sequencing of the *SB* tumors to identify the common insertion sites (CIS); *(ii)* then identified the genes in proximity to the CIS (i.e., the CIS-associated genes); and *(iii)* lastly, determined which CIS-associated genes are differentially expressed in the NEPC versus non-NEPC *SB* tumors (Fig. 1, Step 4). CIS refer to those transposon insertion sites that most frequently occur across multiple independent tumors and therefore are most likely to occur proximal to genes associated with the tumor phenotype (Ranzani et al., 2013; Sarver et al., 2012). Given the volume of transposon insertion junctions that occur randomly throughout the genome, a statistical approach, such as for example Poisson distribution statistics, is needed to separate statistically significant CIS events from random background.

First, we identified SB transposon insertion sites via multiplexed Illumina sequencing of *NPp53-SB(+)* tumors (n = 74), using barcodes to identify individual tumors (Table S6, see Methods). We then determined the statistically relevant insertions across multiple independent tumors by employing the Transposon Annotation Poisson Distribution Association Network Connectivity Environment (TAPDANCE) software (Sarver et al., 2012); to visually depict the data we used a Circos plot (Fig. 5A, Table S7). TAPDANCE statistical analysis identified 122 common insertion sites (CIS), mapping to either the plus or the minus DNA strand, that are well-distributed across the chromosomes throughout the genome. These CIS are shown in the Circos plot with blue and red dots identifying insertion in the minus or plus strand, respectively. Black lines (between the dots and the chromosome numbers) represent the CIS (i.e. areas where there are more insertions than expected based on the null model) (Fig. 5A).

**Figure 5.**
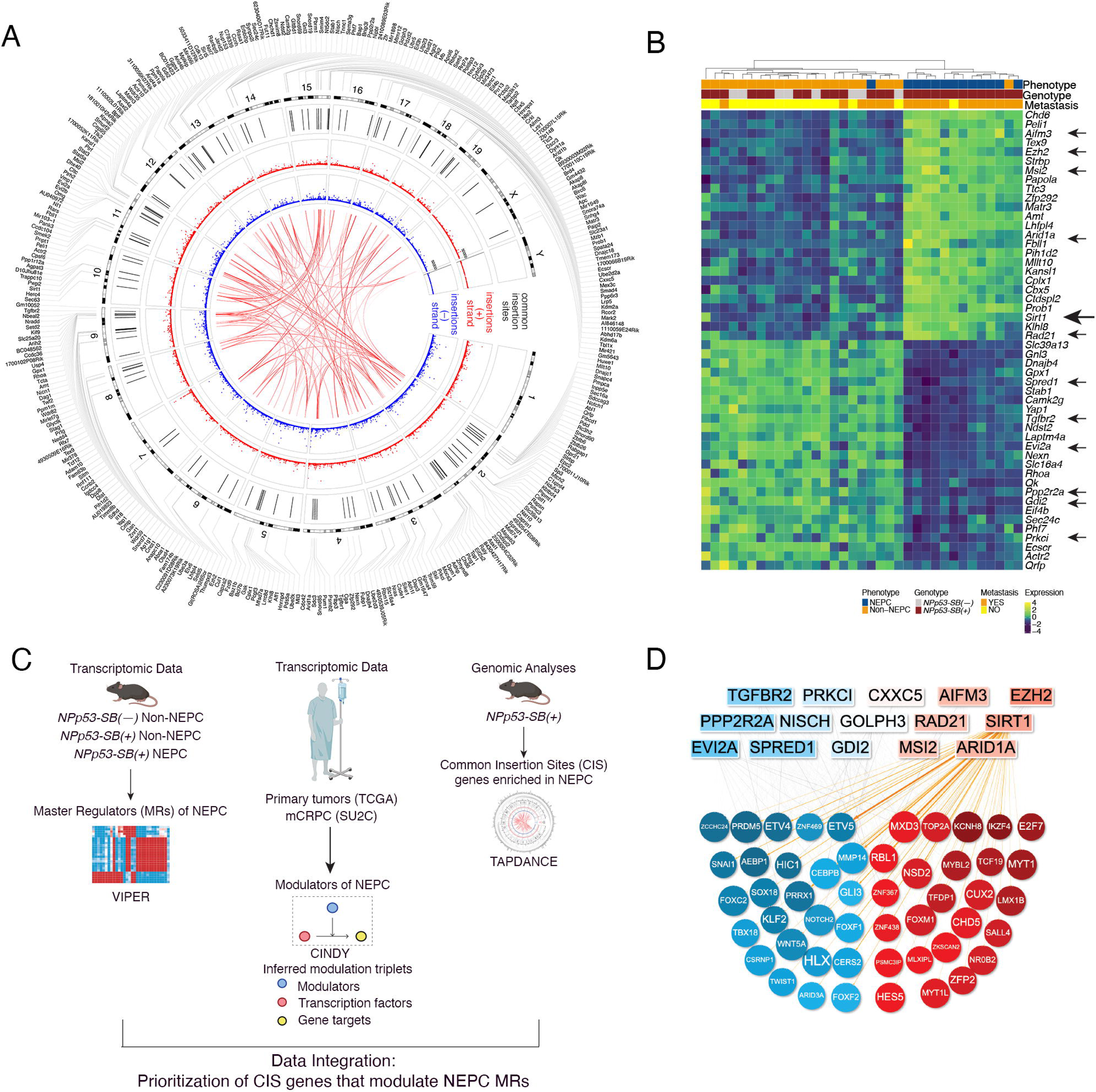
Identification and prioritization of CIS-genes that are mechanistic determinants of NEPC. A. Circos plot of CIS-associated genes identified using the TAPDANCE algorithm. Transposon insertions in the plus (red dots) and minus (blue dots) strands are individually annotated, with corresponding CISs (black lines) and the genes associated to those CISs (outer rim). Red lines in the center of the Circos plot connect CISs that significantly co-occur in tumors. **B.** RNA-seq heatmap showing relative expression levels of the 25 most up-regulated and 25 most down-regulated CIS-genes in NEPC versus non-NEPC *SB* mouse prostate tumors. **C.** Strategy for prioritization of CIS-genes that are mechanistic determinants of NEPC. *i)* The VIPER algorithm was used on the *Sleeping Beauty* cohort to identify MRs specific of the NEPC phenotype (see Fig. 4B); *ii)* The CINDy algorithm was used on the TCGA and SU2C patient datasets to identify proteins that regulate the function of transcription factors (TFs) on target genes (i.e. Modulators); *iii)* we narrowed down the list of Modulators to those that regulate the function of the NEPC MRs, identified from step *i);* and *iv)* we further prioritized CIS-genes that are differentially expressed in NEPC versus non-NEPC *SB* tumors. **D.** Visual representation of the 15 prioritized CIS-genes that are Modulators (top) operating upstream of NEPC MRs (bottom). See also Fig. S4, Tables S6, S7, and S8.

To identify genes proximal to these 122 CIS, we used the TAPDANCE algorithm to select genes within 20 kilobase pairs (kbp) distance from each CIS. Examples of CIS and their location and directionality as related to such genes are shown using the lollipop representation (Fig. S4). One CIS event could not be mapped to any gene within 20kbp, while the remaining 121 CIS events were mapped to 330 CIS-associated genes, depicted as the outer ring of the Circos plot (Fig 5A, Table S7). Furthermore, since TAPDANCE provides both precise CIS mapping coordinates and directionality, we were able to analyze the strandedness of each CIS and to assess CIS co-occurrence across between samples (Fig 5A, red internal lines).

Some CIS events were more frequent in NEPC SB tumors (Table 2, Table S7, see methods), suggesting that these may occur at genes that play a causal role in implementing the NEPC transition. Since most functional CIS events either increase or decrease the expression of the gene mediating their effect, we further prioritized CIS-associated genes based on their differential expression in NEPC vs. non-NEPC *SB* tumors, leading to 75 candidate CIS-associated, NEPC-differentially expressed genes (CIS-genes for short) (Fig. 5B, Table S7).

**Table 2:**
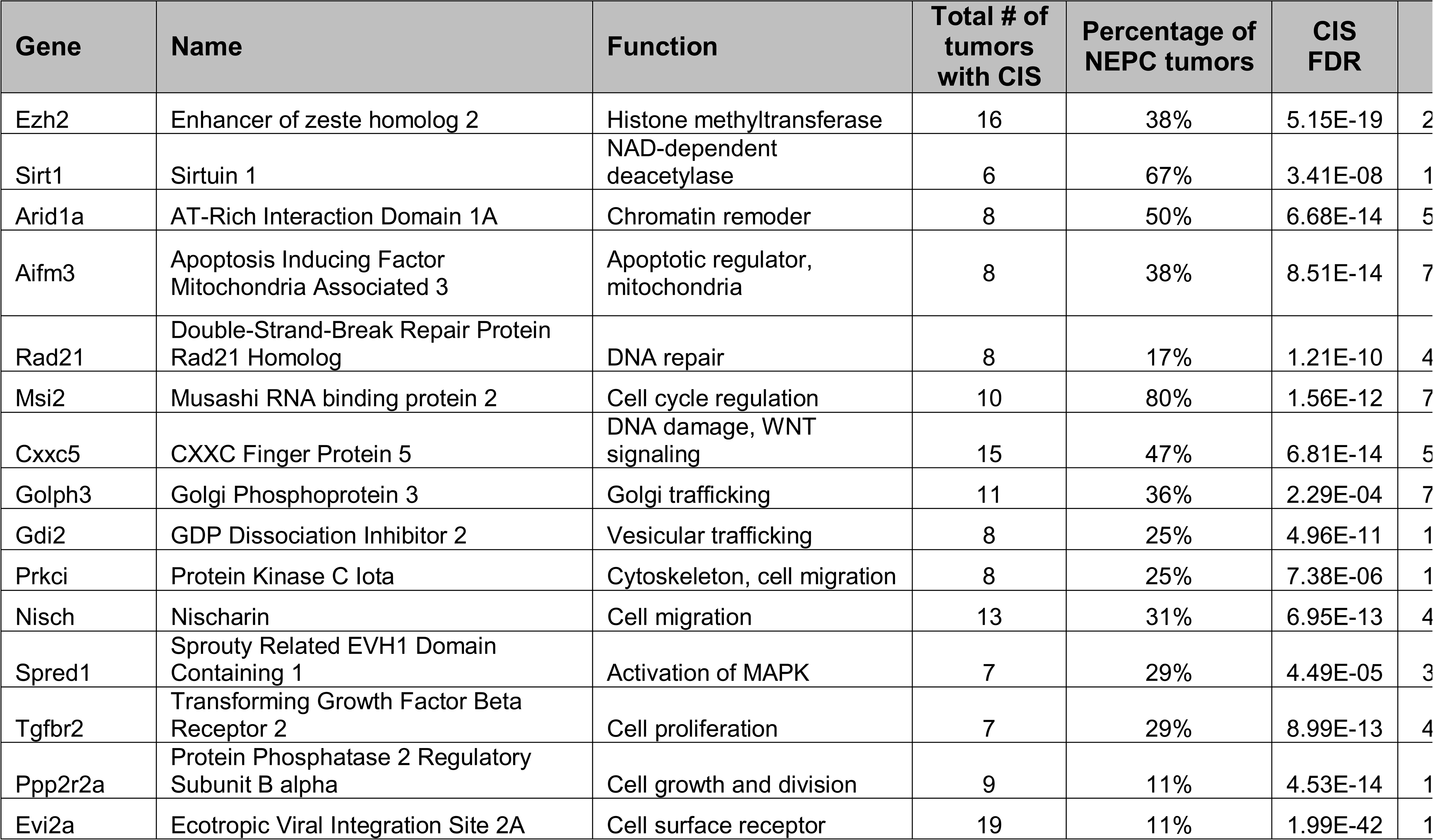
Prioritization of common insertion site (CIS)-genes that are candidate mechanistic determinants of NEPC.

### Prioritization of CIS-associated genes that are mechanistic determinants of NEPC

To prioritize CIS-genes that are most likely to be mechanistic determinants of NEPC (Fig. 1, Step 5), we used a multi-step analysis that integrated transcriptomic and genomic data from the mouse *SB* tumors, and analyses of human NEPC patient cohorts (Fig. 5C,D, Table 2, Table S8). In particular, in previous studies (Chen et al., 2014; Paull et al., 2021) we have shown that causal genetic events affecting tumor cell state occur in genes encoding proteins that can affect the activity of one or more MR proteins (i.e., upstream MR modulators). These can be effectively identified by assessing whether differential abundance of a modulator gene, over a large sample repository (n ≥ 200), can affect the ability of an MR protein to regulate its targets. This is assessed by determining the statistical significance of the conditional mutual information (CMI) <u>![MR; T|M]</u>, where MR is a Master Regulator, T represents the genes identified as its transcriptional targets and M is the candidate modulator protein. The CMI can be effectively assessed by analyzing RNA-seq profiles using CINDy (Giorgi et al., 2014), the most recent implementation of the Modulator Inference by Network Dynamics (MINDy) algorithm (Wang et al., 2009a). We thus hypothesized that the CIS representing causal, mechanistic determinants of NEPC cell state would be proximal to genes identified as CINDy modulators of one or more NEPC MRs.

We thus integrated the following analyses (Fig. 5C): *(i)* Candidate MRs of the NEPC vs. non-NEPC tumor state transition, which were identified using the VIPER algorithm (Alvarez et al., 2016) (from Fig. 4B). Among ∼2,500 genes encoding for TF and coTF proteins, only those producing a VIPER-based abs(NES) 2: 9 were labeled as candidate MRs. *(ii)* We then used CINDy to prioritize candidate MR activity modulators among ∼3,800 signaling proteins, as in (Giorgi et al., 2014). We performed this analysis on two independent patient cohorts, namely *(i)* the Cancer Genome Atlas (TCGA) cohort, which is comprised of primary prostate tumors that had not undergone treatment (N=498, (Cancer Genome Atlas Research, 2015)), and *(ii)* the SU2C mCRPC cohort introduced above, (N=266, (Abida et al., 2019)). Briefly, for each cMR andidate modulator gene [M], and MR transcriptional target [T]—as inferred by the ARACNe algorithm (Basso et al., 2005)—we computed the statistical significance of the conditional mutual information ![cMR; T|M], using RNA-seq profiles from both human cohorts, independently. The statistical significance (p-value) of each candidate modulator gene was assessed by integrating the p-values across all of its targets, as described in (Giorgi et al., 2014), and across cohorts. Statistical significance was assessed at p ≤ 0.05 (FDR corrected).

To prioritize candidate CIS-genes that are mechanistic determinants of NEPC, we assessed each candidate by integrating three independent p-values: *(i)* the CIS-gene p-value as assessed by TAPDANCE; *(ii)* the p-value of its differential expression in NEPC vs. non-NEPC samples, as shown in Fig. 5B; and *(iii)* the p-value of its CINDy-inferred activity as an NEPC MR modulator (Table 2; Table S8). P-values from these analyses were integrated using Fisher’s method and used to rank all CIS-genes; genes not reported by any of the analyses were assigned p-value of 1.

The analysis identified 15 CIS-genes as high-likelihood candidate drivers of the NEPC tumor state (Fig. 5D, Table 2, Table S8). Among these, some have previously been shown to be involved in NEPC, such as the lysine methyltransferase, Ezh2 (for Enhancer Of Zeste 2 Polycomb Repressive Complex 2 Subunit) (Dardenne et al., 2016; Ku et al., 2017). Other candidates, such as Tgfβ type II receptor (Tgfbr2), have not been previously directly implicated in NEPC, but are associated with pathways (in this case TGFβ signaling) known to be involved in plasticity and prostate cancer progression (Hao et al., 2018). Other candidates, such as the Musashi RNA binding protein 2, Msi2, have not been previously implicated in NEPC, but have notably functions in cancer and cell stemness (Kharas et al., 2010).

### SIRT1 promotes NEPC in human prostate cancer

Among these candidates, we focused our subsequent analysis the nicotinamide adenosine dinucleotide (NAD)-dependent deacetylase *sirtuin 1* (SirT1). Notably, Sirt1 had the highest VIPER measured differential protein activity in NEPC vs. non-NEPC tumors, an indicator of its predicted functional role in NEPC, and was also associated with the CIS that occurred in the highest percentage (67%) of NEPC *SB* tumors (Table 2). Additionally, *SIRT1* has been shown to be up-regulated in prostate cancer (Huang et al., 2021; Huffman et al., 2007), including in NEPC contexts (Natani et al., 2021; Ruan et al., 2018). Notably, our integrative approach not only places SIRT1 as a candidate top that regulates NEPC, but it also shows that its regulon of downstream factors includes several known NEPC MRs, including MYT1, NEUROD1, INSM1 and ASCL1 (Fig. 6A). Furthermore, SIRT1 has been shown to be targetable using small molecule inhibitors that block its lysine deacetylase activity (Adams et al., 2024; Napper et al., 2005; Westerberg et al., 2015). Thus, we performed validation studies to evaluate the functional relevance of SIRT1 for NEPC in human prostate cancer cell lines (Fig. 1, Step 6).

**Figure 6.**
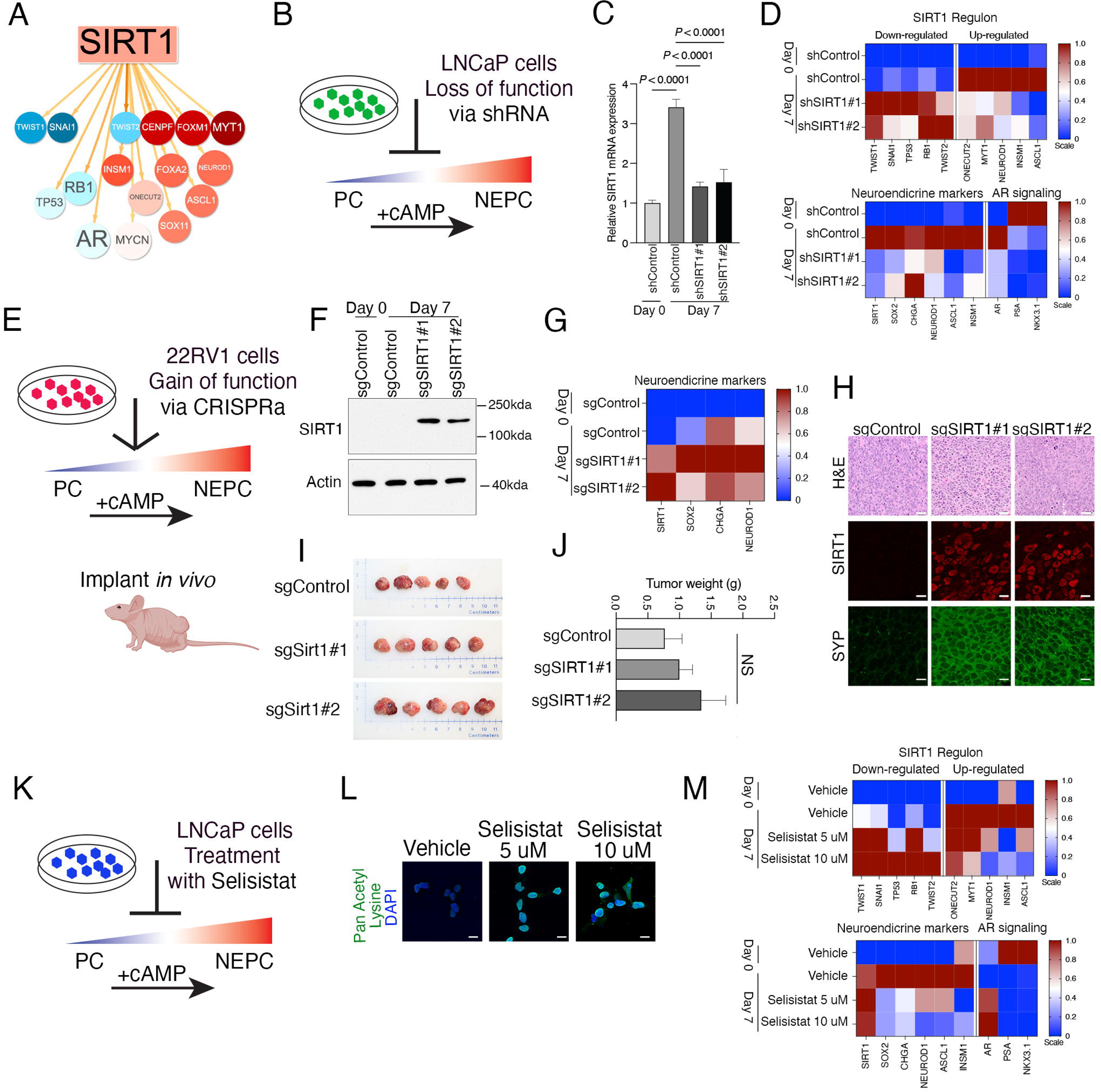
SIRT1 promotes NEPC in human prostate cancer cell models. **A**. Visual depiction of a subset of the SIRT1 regulon showing NEPC MRs that are predicted to be up-regulated (red) or down-regulated (blue) by SIRT1 in prostate cancer. **B-D**. <u>Loss-of-function of SIRT1 in LNCaP cells</u>. **B**. Schematic representation showing that LNCaP cells can be induced to transition from prostate cancer (PC) to NEPC by treatment with db-cAMP and IBMX (abbreviated cAMP). Silencing SIRT1 is predicted to inhibit this transition. **C.** Relative expression of SIRT1 **D**. Heatmap representation of relative expression levels of the SIRT1 regulon (top) and known markers of NEPC or androgen signaling (bottom). **E-J**. <u>Gain-of-function of SIRT1 in 22RV1 cells</u>. **E**. Schematic representation showing treatment of 22RV1 cells with db-cAMP and IBMX (abbreviated cAMP). SIRT1 expression was induced with CRISPRa. Studies were done in vitro (F,G) and in vivo (H-J). **F**. Western blot showing protein expression levels of SIRT1 in the 22RV1 cells. Actin, control for protein loading; position of molecular weight markers are shown. **G**. Heatmap representation of relative gene expression levels of known markers of NEPC. **H-J**. Results of 22RV1 cells grown orthotopically in host *nude* mice. **H.** Representative images of the 22RV1 prostate tumors. Shown are hematoxylin and eosin (H&E) staining, immunofluorescence (IF) staining for SIRT1 and Synaptophysin (SYP). Scale bars represent 20 µm for H&E, and 10 µm for IF images. **I**. Prostate tumors collected at the time of sacrifice. **J**. Summary of tumor weights. **K-M**. <u>Pharmacological inhibition of SIRT1 in LNCaP cells</u>. **K**. Schematic representation showing that LNCaP cells can be induced to transition from prostate cancer (PC) to NEPC by treatment with db-cAMP and IBMX (abbreviated cAMP). Treatment with the SIRT1 inhibitor Selisistat is predicted to inhibit this transition. **L**. Relative levels of lysine acetylation following treatment of LNCaP cells with 5 or 10 µM Selisistat. Scale bars represent 20 µm. **M**. Heatmap representation of relative expression levels of the SIRT1 regulon (top) and known markers of NEPC or androgen signaling (bottom). See also Fig. S5.

To evaluate the functions of SIRT1 in NEPC, we performed studies in LNCaP cells (Horoszewicz et al., 1980), which are an AR-dependent prostate cancer cell line that can be induced to differentiate to a neuroendocrine-like phenotype when deprived of androgens and treated with agents that increase the intracellular levels of cyclic AMP (cAMP), such as dibutyryl-cAMP (db-cAMP) and 3-isobutyl-1-methylxanthine (IBMX, hereafter we refer to this as NEPC induction and abbreviate pictorially as +cAMP) (Fig. 6B) (Bang et al., 1994; Burchardt et al., 1999; Shen et al., 1997). We found that SIRT1 is expressed at low levels prior to NEPC induction (day 0), while its expression is significantly increased following NEPC induction (day 7); this increased expression of SIRT1 was abrogated using 2 independent SIRT1 shRNA (Fig. 6B,C). Notably, NEPC induction of the control LNCaP cells resulted in an increase in expression of the genes comprising the SIRT1 regulon that are predicted to be activated by SIRT1 (ONECUT2, MYT1, NEUROD1, INSM1 and ASCL1), and a corresponding decrease in expression of those predicted to be down-regulated by SIRT1 (TWIST1, SNAI1, TP53, RB1 and TWIST2) (Fig. 6D, top). However, this change in gene expression was abrogated by depletion of SIRT1 (Fig. 6D, top). Furthermore, NEPC induction of the LNCaP cells (day 7) resulted in an increased expression of NEPC markers (SOX2, Chromogranin A (CHGA), NEUROD1, ASCL1 and INSM1), which was abrogated by depletion of SIRT1 (Fig. 6D, bottom), supporting the idea that SIRT1 is necessary for the NEPC differentiation of LNCaP cells.

To extend these findings, we performed studies using a second human prostate cancer cell line, 22RV1 (Sramkoski et al., 1999) that has not been extensively used to study neuroendocrine-related phenotypes. We found that SIRT1 expression was similarly low in both the untreated cells (day zero) as well as following NEPC induction (day 7) (Fig. 6F-G, Fig. S5A). Therefore, we reasoned that 22RV1 cells represented an ideal model to investigate the consequences of gain-of-function of SIRT1 for NEPC phenotypes *in vitro* as well as *in vivo* following orthotopic growth in host mice (Fig. 6E).

To activate expression of SIRT1 in the 22RV1 cells, we used a CRISPR activation (CRISPRa) model previously described (Arriaga et al., 2024). We engineered the 22RV1 CRISPRa cells to express a control single guide RNA (sgRNA) or 2 independent sgRNAs for SIRT1. Both SIRT1sgRNAs resulted in a robust increase in expression of SIRT1 protein and mRNA (Fig. 6F-G, S5A). Notably, while expression of SIRT1 had a modest or negligible effect on cell proliferation and colony formation of the 22RV1 cells (Fig. S5B-D), it resulted in a striking increase in the expression of relevant NEPC markers, including SOX2, CHGA, and NEUROD1 (Fig. 6G). Even more striking, orthotopic implantation of these 22RV1 CRISPRa cells into the prostate of male host mice revealed a modest effect on tumor growth (Fig. 6I,J), but a striking effect on the expression of markers of NEPC, including Synaptophysin (SYP) and INSM1, even though it had a modest effect on AR expression (Fig. 6H, Fig. S5E). These findings suggest that SIRT1 promotes an NEPC phenotype, while not having an overall effect on prostate tumorigenicity.

Lastly, we asked whether pharmacological inhibition of SIRT1 could inhibit NEPC features in human prostate cancer cells (Fig. 6K). For these studies, we used the SIRT1-specific small molecule inhibitor Selisistat, which has been demonstrated to inhibit the lysine deacetylase activity of SIRT1 in other cellular contexts (Adams et al., 2024; Napper et al., 2005; Westerberg et al., 2015). In the current studies, we found that lysine deacetylase activity, but not overall cellular viability, was effectively inhibited at 5 or 10 μM of Selisistat in the LNCaP (Fig. 6L), which is similar to doses used in other cell models (Adams et al., 2024). Furthermore, pharmacological inhibition of SIRT1 abrogated the observed NEPC-induced LNCaP changes in gene expression of the SIRT1 regulon, such as ONECUT2 and MYT1 (Fig. 6M, top), as well as neuroendocrine markers, including SOX2, INSM1, and ASCL1 (Fig. 6M, bottom). Taken together, these findings suggest that SIRT1 promotes NEPC features in human prostate cancer cells, while its depletion or pharmacological inhibition abrogates the NEPC phenotype in these contexts.

## Discussion

Over the past several decades, forward genetic screens have become powerful tools for identifying the function of genes and pathways, providing valuable insights into the complexity of cancer phenotypes. Importantly, forward genetic screens can identify genes or pathways that may be targets for therapeutic intervention, particularly in the context of disease-related phenotypes. In this regard, *Sleeping Beauty* mutagenesis has proven to be a valuable approach for performing unbiased screens of cancer drivers (Beckmann and Largaespada, 2020; Copeland and Jenkins, 2010). A particular benefit of the *Sleeping Beauty* approach is that mutagenesis occurs in autochthonous mice and therefore arises somatically during disease progression in the context of the native tumor environment, analogous to the accumulation of mutations during cancer progression in humans. In the current study, we have used a *Sleeping Beauty* mutagenesis screen to investigate molecular drivers of advanced prostate cancer. Furthermore, we have interpreted the screening results using an integrative approach that encompasses analyses of the tumor phenotype with analyses of transcriptomic data and genomic data of transposon insertion sites. These studies have led to the identification of the NAD deacetylase SIRT1 as a targetable driver of NEPC and introduce a generalizable approach for the interpreting findings of unbiased *in vivo* mutagenesis screens.

Unlike previous *Sleeping Beauty* screens that have been implemented in prostate cancer mouse models based on loss-of-function of the *Pten* tumor suppressor gene (Ahmad et al., 2016; Rahrmann et al., 2009), our study utilized a mouse model with loss-of-function of both *Pten* and *Trp53* (*NPp53* mice) (Zou et al., 2017). Basing the screen on the *NPp53* mice allowed us to identify drivers of advanced prostate cancer and, in particular, to study molecular determinants of NEPC, which is a lethal form of the disease that has emerged as a consequence of disease resistance (Beltran et al., 2019; Watson et al., 2015). Notably, the NEPC phenotypes in the *NPp53-SB(+)* mice arise in the context of intact androgen signaling and in the absence of any drug treatment; therefore, these findings provide a unique opportunity to study the *de novo* cellular events that give rise to lethal prostate cancer.

A second innovation of our study is the introduction of a computational pipeline that using state-of-the-art algorithms to leverage a comprehensive analysis of the phenotypic, transcriptomic, and genomic analyses of the mouse *SB* tumors with integration to human prostate cancer patient cohorts to identify and prioritize candidate CIS genes that are mechanistic determinants of NEPC. Indeed, a significant challenge of *SB* and other genetic screens is the prioritization of candidate drivers of the cancer phenotype. Our study highlights the value of integrating genomic and transcriptomics data to gain insights that facilitate the selection of key genes most likely to drive the phenotype of interest. Thus, we have combined transcriptomic and genomic data from the *SB* mice with algorithms to reverse engineer regulatory networks for both mouse and human prostate cancer, which has allowed us to identify CIS genes that are most likely to that modulate relevant transcriptional programs within the context of our disease model. Our current study builds upon our previous work, which has shown the value of using transcriptomics data to reverse engineer gene regulatory networks, not only to identify interactions between transcription factors (TFs) and target genes, but ultimately to unravel genes that modulate the activity of those TFs on their target genes (*i.e.* Modulators) (Giorgi et al., 2014; Paull et al., 2021; Vasciaveo et al., 2023).

Among the top candidate genes that were identified using this prioritization approach, SIRT1 emerged as a lead candidate that is a mechanistic determinant of NEPC. SIRT1 is a member of a family of genes that were identified in non-mammalian contexts based on their roles in extending lifespan (Haigis and Sinclair, 2010), and is now known to be an NAD lysine deacetylase that facilitates the formation of heterochromatin (Vaquero et al., 2004). SIRT1 has been shown to play an important role in the nervous system, as well as in neurogenerative disease and neuroendocrine disease (Fujita and Yamashita, 2018). It is also a known regulator of cancer, particularly in conjunction with p53, however, whether it functions as an oncogene or tumor suppressor appears to be context-dependent (Lin and Fang, 2013).

A role for SIRT1 in prostate cancer has been proposed for some time, however, like other contexts, whether or not it functions has a positive or negative impact on prostate cancer has been somewhat unclear. Previous studies have shown that its loss results in preinvasive cancer phenotypes, whereas other studies have shown that promotes prostate cancer progression. Our studies serve to reconcile these findings since we show that SIRT1 does not affect tumor growth in vivo or cell proliferation in vitro, rather it promotes an NEPC phenotype. Notably, SIRT1 was originally identified in yeast as a regulator of the SWI/SNF chromatin complex, and thus it is tempting to speculate that the functions of SIRT1 functions in NEPC in prostate cancer are mediated by its effects on the SWI/SNF complex (Mittal and Roberts, 2020) particularly in light of the key role of the SWI/SNF in prostate cancer (Cyrta et al., 2020).

Our studies further suggest that SIRT1 targeting may invert progression to NEPC. SIRT1 drugs like Selisistat are FDA-approved and have been previously tested in Phase I trials, thus being accepted in the clinic (Westerberg et al., 2015). This makes SIRT1 an attractive target for intervention for lethal prostate cancer, which can be further evaluated in future clinical trials.

## Materials and Methods

### Generation and analysis of Sleeping beauty *(SB)* prostate cancer mouse models

All experiments using animals were performed according to protocols approved by the Institutional Animal Care and Use Committee (IACUC) at Columbia University Irving Medical Center. Mice were housed in pathogen-free barrier conditions under 12-h light/dark cycles and with temperature and humidity at 20–25 °C and 30–70%, respectively. Genetically-engineered mouse models (GEMMs) in this study utilize the *Nkx3.1*^CreERT2/+^ allele to activate an inducible Cre recombinase in the prostatic epithelium (Wang et al., 2009b). The mice used for the Sleeping beauty mutagenesis are the Cre*-* inducible transposase allele (*Rosa26-LSL-SB11* (Geurts et al., 2003; Starr et al., 2009)) with or without the *T2/Onc2* allele (NCI Mouse Repository, Frederick National Laboratory, strain 01XGA) (Dupuy et al., 2005); together these comprise the *SB* mice wherein the *SB(–)* have only the *Rosa26-LSL-SB11* allele while the *SB(+)* mice have both the only the *Rosa26-LSL-SB11* and the*T2/Onc2* alleles. Since the *T2/Onc2* transposon is located on chromosome 1, insertions on this chromosome were disregarded in subsequent analysis to overcome complications related to local hopping (Liang et al., 2009). The *SB(–)* and *SB(+)* mice were crossed with the *NPp53* mice (*Nkx3.1^CreERT2/+^; Pten^flox/flox^; Trp53^flox/flox^* (Zou et al., 2017)) to generate the *NPp53-SB(–)* mice *(Nkx3.1^CreERT2^; Pten^flox/flox^; Trp53^flox/flox^; R26-LSL-SB11^flox/flox^)* and the *NPp53-SB(+)* mice *(Nkx3.1^CreERT2^; Pten^flox/flox^; Trp53^flox/flox^; R26-LSL-SB11^flox/flox^; T2/Onc2*). Alternatively, the *SB(–)*and *SB(+)* mice were crossed with the *NP* mice (*Nkx3.1^CreERT2/+^; Pten^flox/flox^* (Floc’h et al., 2012)) to generate the *NP-SB(–)* mice *(Nkx3.1^CreERT2^; Pten^flox/flox^; R26-LSL-SB11^flox/flox^)* and the *NP-SB(+)* mice *(Nkx3.1^CreERT2^; Pten^flox/flox^; R26-LSL-SB11^flox/flox^; T2/Onc2)*. A summary of all mice used in this study is provided in Table S1.

All studies were performed using littermates that were genotyped prior to tumor induction. Since our focus is on prostate cancer, only male mice were used. Mice were induced at 2-3 months of age by administration of tamoxifen (Sigma-Aldrich T5648, St. Louis, MO, USA) using 100 mg/kg (in corn oil) once daily for 4 consecutive days. Following tamoxifen-induction, mice were monitored three times weekly, and euthanized when their body condition score was <1.5, or when they experienced body weight loss ≥ 20% or signs of distress, such as difficulty breathing or bladder obstruction. At their endpoint, mice were euthanized via carbon dioxide inhalation followed by cervical dislocation and were necropsied. Harvested tissues were visualized using an Olympus SZX16 microscope. Tissues were snap-frozen in liquid nitrogen for DNA or RNA isolation, or were fixed in 10% formalin (Fisher Scientific, Fair Lawn, NJ, USA) for hematoxylin and eosin (H&E), immunohistochemical and immunofluorescence staining.

Histopathological scoring of the prostate tumors was assessed independently by two pathologists (SdB and AR) based on evaluation of H&E-stained tissues, some with confirmation by immunohistochemistry for synaptophysin and Ki67; analyses are summarized in Table S1. Histomorphologic features indicative of neuroendocrine differentiation include: *(i)* organoid nesting trabecular growth pattern, with peripheral palisading, and rosette formation; *(ii)* tumor cells smaller than the diameter of 3 small lymphocytes, have scant cytoplasm, with oval or spindled hyperchromatic nuclei; and *(iii)* mitotic count is high and extensive necrosis is frequently present (Ittmann et al., 2013; Shappell et al., 2004).

Immunostaining was done using 3 μm formalin-fixed sections as described (Zou et al., 2017). Briefly, sections were deparaffinized in xylene followed by heat-induced antigen retrieval using Citrate-based Antigen Unmasking Solution at pH 7.0 (Vector Labs, Newark, CA, USA). Sections were blocked in 10% normal goat serum, incubated with primary antibodies overnight at 4 °C, and with secondary antibodies for 1h at room temperature. For immunohistochemistry, signal detection and visualization were performed using the Vectastain ABC system followed by NovaRed Substrate Kit (Vector Labs, Newark, CA, USA), counterstained with Hematoxylin and mounted with Clearmount (American Master*Tech Scientific). Images were captured using an Olympus VS120 whole-slide scanning microscope. For immunofluorescence, tissues were incubated with primary and secondary antibodies as above and then stained with DAPI and mounted with Vectashield antifade mounting medium (Vector Labs, Newark, CA, USA). Images were captured using a Leica TCS SP5 confocal microscope. Quantification of Ki67 staining was performed manually on a minimum of 8,000 cells per group based on 3 independent tumors with a minimum of 5 sections per tumor, as described in (Zou et al., 2017). All antibodies and secondary antibodies are described in Key Resources Table.

Validation of mRNA expression levels was done by quantitative real time PCR using the QuantiTect SYBR Green PCR kit (Qiagen, Germantown, MD, USA) using mouse *GADPH* as the control (Zou et al., 2017). Primer sequences are provided in the Key Resources Table.

### Transcriptomic of Sleeping beauty *(SB)* prostate cancer mouse models

RNA sequencing was performed as described (Vasciaveo et al., 2023) on a total of 35 RNA samples from 32 independent mouse prostate tumors, including 8 *NPp53-SB(–)* tumor samples and 27 *NPp53-SB(+)* tumor samples, of which 13 were histologically characterized as NEPC and 22 as non-NEPC (Table S1). We also analyzed 4 *NP-SB(–)* and 4 *NP-SB(+)* tumors, all of which were non-NEPC (Table S1). RNA was prepared from snap-frozen tissues that were homogenized in TRIzol (ThermoFisher Scientific, Waltham, MA, USA) and extracted using the MagMAX-96 total RNA isolation kit (ThermoFisher Scientific, Waltham, MA, USA). Total RNA was enriched for mRNA using poly-A pull-down; only samples having between 200 ng and 1 µg and with an RNA integrity number (RIN) > 8 were used.

Libraries were made using an Illumina TruSeq RNA prep-kit v2 or TruSeq Stranded mRNA library prep kit and sequenced using an Illumina HiSeq2500/4000 or NovaSeq6000. RNA-seq profiles were generated by mapping mRNA reads to the mouse reference genome (version GRCm38 mm10), using *kallisto* v0.44.0. RNAseq raw counts were normalized and variance stabilized using DESeq2 (v.1.36.0) package (Bioconductor) in R-studio 2023.03.0+385, R v.4.2.3 (R Foundation for Statistical Computing).

For the generation of gene expression heatmaps, variance stabilized counts were z-scaled by row and visualized using the *ComplexHeatmap* (v2.14.0) package (Bioconductor) in R-studio 2023.03.0+385, R v.4.2.3 (R Foundation for Statistical Computing). Principal Component Analysis (PCA) was performed using the plotPCA function from the *DESeq2* (v1.42.1) package, using the top 500 highly variable genes computed from variance stabilized count data. To compare with the mouse RNA sequencing data with human NEPC gene signatures, we used the AR, NEURO I and NEURO II gene expression sets published (Labrecque et al., 2019). Data were visualizing and plotted as a heatmap.

To identify genes differentially expressed in the histologically-characterized NEPC vs non-NEPC SB mouse tumors, we used *edgeR* (v4.0.16). Read counts were modeled using Genewise Negative Binomial Generalized Linear Models with Quasi-Dispersion Estimation (glmQLFit) after having estimated gene dispersion using Empirical Bayes Tagwise Dispersions for Negative Binomial GLMs (estimateGLMTagwiseDisp). The contrast matrix was generated by comparing SB(+) tumors that displayed NEPC with those that did not display NEPC. The list of statistically significantly (FDR < 0.1, Benjamini-Hochberg corrected) differentially expressed genes is provided in Table S2; the dataset is provided in GSE271053.

### Protein activity-based cluster analysis

These analyses were done using the RNA sequencing data from the SB tumors including the 13 histologically-characterized NEPC tumors and the 22 histologically-characterized non NEPC tumors (see above). First, a differential gene expression signature (DGES) was computed for each sample (Table S2). Then the VIPER algorithm (Alvarez et al., 2016) was used to transform each DGES into a differential protein activity signature (DPAS). A summary of VIPER protein activity is provided in Table S3. Cluster analysis was performed using the hierarchical clustering algorithm as implemented using *hclust* (v4.3.2), with Ward’s agglomeration method (Fionn and Legendre, 2014). Spearman’s correlation of the respective protein activity profiles was used as a sample-to-sample distance metric. For visualization purposes, only the 15 most statistically significant candidate MRs are shown, as assessed by p-value integration across the samples of each cluster (n = 3), using Stouffer’s method. Relative activity of each MR was then visualized across all samples. A summary of protein activity of each cluster is provided in Table S4.

### OncoMatch protein activity-based analysis of mouse and human tumors

This analysis comprises three steps. First, DGES was computed for each mouse and human sample (Step 1, Table S2). Then the VIPER algorithm (Alvarez et al., 2016) was used to transform each DGES into a differential protein activity signature (DPAS) (Step 2, Table S3). Finally, the most differentially active proteins (both most activated and most inactivated) were selected as candidate Master Regulator (MR for short) proteins of each individual patient-derived tumor sample from relevant human cohorts and used to assess the fidelity of each mouse tumor, using the OncoMatch algorithm (Vasciaveo et al., 2023) (Step 3, Table S4).

#### Step 1 – Differential Gene Expression Signature generation

We independently generated a DGES for each mouse and patient-derived tumor sample by computing Z-scores obtained by dividing the difference between the expression of each gene in the sample and the mean expression in the associated cohort by the standard deviation. The mouse cohort comprises 35 tumors including the 13 histologically-characterized NEPC tumors and 22 histologically-characterized non NEPC tumors (Table S1). Human tumor profiles were collected from publicly available databases, including: (1) The Beltran cohort (n=49 patients), comprising 34 adenocarcinoma and 15 NEPC patients, as downloaded on November 12, 2020 from cBioPortal as FPKM measurements (Beltran et al., 2016) and (2) The SU2C cohort (n=266 patients), comprising 210 non-NEPC, 22 NEPC patients and 34 unclassified tumors, as downloaded on October 4, 2019 from cBioPortal as FPKM measurements based on the 2019 SU2C-mCRPC dataset (Abida et al., 2019).

#### Step 2 – Differential Protein Activity Signature generation and candidate MR identification

Differentially active proteins were identified by analyzing each DGES from Step 1, using the VIPER (Alvarez et al., 2016) and metaVIPER (Ding et al., 2018) algorithms, with the species-appropriate regulatory networks, described in (Vasciaveo et al., 2023). The GEMM signatures were processed by metaVIPER by integrating both the human SU2C network and a GEMM network as in (Vasciaveo et al., 2023). The protein activity signatures are provided in Table S3.

#### Step 3 – OncoMatch analysis of SB prostate tumors and human prostate cancer patients

To evaluate the fidelity (i.e., similarity) of *SB* mouse prostate tumors to human prostate cancer patients we leveraged the OncoMatch algorithm (Vasciaveo et al., 2023). Briefly, OncoMatch measures the enrichment of the top Master Regulator proteins of a human tumor—as assessed by VIPER/metaVIPER analysis—in proteins differentially active in a tumor model (i.e., in this case the mouse *SB* tumors). For this purpose, we selected the 50 most differentially active proteins in each human tumor—including the 25 most activated (25↑) and 25 most differentially inactivated proteins (25↓), as assessed by VIPER/metaVIPER—as candidate Master Regulator proteins (MRs for short). Indeed, we have shown that 25 to 50 MRs can account for >80% of a tumor somatic mutations in their upstream pathways across all of TCGA (Paull et al., 2021) and have shown that this number of MRs supports effective assessment of model fidelity (Vasciaveo et al., 2023).

The enrichment of patient tumor-specific MRs in proteins differentially active in the GEMM tumors was assessed using the aREA algorithm, as also described in (Vasciaveo et al., 2023). For this purpose, murine proteins were *humanized* (*i.e.*, mapped to their human protein orthologs) before running the aREA algorithm as shown in (Vasciaveo et al., 2023). Normalized Enrichment Scores (NES) were converted to *p*-values and the Bonferroni method was used to correct for multiple hypothesis testing. The value of OM_Score_ = -Log_lO_ (p - value) > 5 was used as an MR conservation threshold to assess effective tumor/model fidelity. The OncoMatch data is provided in Table S5.

### Identification of common insertion sites (CIS)

Targeted genomic sequencing was performed to identify transposon insertion sites in *NPp53-SB(+)* prostate tumors following the protocol published in (Janik and Starr, 2013). We analyzed 74 tumors for which the *NPp53-SB(+) T2/Onc2* transposition was confirmed by PCR (Fig. S1A). Genomic DNA was extracted and digested with restriction enzymes flanking the left (BfaI) or right (NlaIII) side of the transposon, as in (Janik and Starr, 2013). The resulting 148 samples were subjected to linker-mediated-PCR (LM-PCR) as published in (Janik and Starr, 2013). Samples were prepared using the primers in the Key Resources Table and the barcodes described in Table S6. Samples were sequenced using the Illumina HiSeq with 100bp read (Azenta, Burlington, MA, USA).

Common insertion sites (CIS), which are defined as regions of the genome having transposon insertions (i.e., integration sites) across multiple independent tumors at a significantly greater frequency than that expected by chance (Dupuy et al., 2005), were identified using the Transposon Annotation Poisson Distribution Association Network Connectivity Environment (TAPDANCE) software (v3.1), which both maps insertion reads and computes statistically significant CIS (Sarver et al., 2012). Reads were mapped using the University California Santa Cruz indexed bowtie 1.2.3 compatible mm10 reference genome. All default parameters were used, and the $library percent, which discards insertion regions that are represented by a low number of reads, was set to .001 (i.e., insertion regions represented by lower than .001 percent of total reads assigned to the tumor are discarded).

This analysis led to the identification of 122 unique CIS. For 121 of the 122 CIS, we identified 330 CIS-associated genes based on annotated genomic location within 20 kbp of the CIS. One CIS lacked any annotated gene feature within 20 kbp of the insertion. Table S7 provides a list of the 330 CIS-associated genes together with the standardized TAPDANCE output. For each CIS-associated gene we also report the proportion of NEPC tumors calculated based on two-sided binomial test (*stats*) using p = proportion of neuroendocrine samples (27/74) for neuroendocrine vs adenocarcinoma.

The 330 CIS-associated genes were represented in a Circos plot, using the *circlize* (0.4.16) package. Strandedness was assessed from the raw TAPDANCE mapping output file, with the height of the dots representing each transposon’s read count. CIS were marked based on TAPDANCE output file (Table S7) and gene positions were acquired by *bioMart* (2.60.1) and plotted in the outer side of the Circos plot. Co-occurrence analysis of pairs of CIS-associated genes was performed using the somaticInteractions function using the *maftools* (2.20.0) package and is represented in the center of the Circos plot.

Lollipop plots for selected CIS-associated genes were generated using *trackViewer* (1.41.5) and *GenomicRanges* (1.57.1) packages. Gene structures were obtained from the mm10 assembly, using the *TxDb.Mmusculus.UCSC.mm10.knownGene* (3.10.0) package. Transposon insertion data from TAPDANCE output were plotted, with insertion heights representing read counts. Insertions in the same direction as gene transcription are shown with red arrows, opposite in green.

### Prioritization of CIS genes representing candidate NEPC regulators

We prioritized CIS genes that promote NEPC as follows; all data is summarized in Table S8:

#### Step 1 – NEPC MR inference

The NEPC vs. non-NEPC differential gene expression signature (NEPC DGES) was computed as described above. Candidate MRs of the NEPC vs. non-NEPC tumor state transition were identified by analyzing the NEPC DGES using the VIPER algorithm (Alvarez et al., 2016), based on a GEMM-derived regulatory network (Vasciaveo et al., 2023). Among ∼2,500 genes encoding for TF and coTF proteins, only those producing a VIPER-based abs(NES) 2: 9 were labeled as candidate MRs.

#### Step 2 – CINDy-based identification of proteins representing MR upstream modulators

We then used CINDy to prioritize candidate modulators of MR activity among ∼3,800 signaling proteins, as in (Giorgi et al., 2014), based on the analysis of RNA-seq profiles from human primary prostate tumors from the Cancer Genome Atlas (TCGA) patient cohort (N=498 (Cancer Genome Atlas Research, 2015)) and metastatic CRPC biopsies from the SU2C patient cohort (N=266, (Abida et al., 2019)). Briefly, for each cMR from Step 2, candidate modulator gene [M], and MR transcriptional target [T]—as inferred by the ARACNe algorithm (Basso et al., 2005)—we computed the statistical significance of the conditional mutual information ![cMR; T|M], using RNA-seq profiles from both human cohorts, independently. The statistical significance (p-value) of each candidate modulator gene was assessed by integrating the p-values across all of its targets, as described in (Giorgi et al., 2014), and across cohorts. Statistical significance was assessed at p ::: 0.05 (FDR corrected).

#### Step 3 – Integrative Analysis

To prioritize candidate CIS-associated genes representing mechanistic NEPC determinants, we assessed each candidate by integrating three independent statistics, including: (1) its p-value as assessed by TAPDANCE analysis, (2) its differential expression in NEPC vs. non-NEPC samples, and (3) their CINDy-predicted activity as NEPC MR modulators. The p-values generated for each of these three analyses was integrated using Fisher’s method. The data are presented in Table S8.

### Functional analysis in cell-based models

LNCaP cells (ATCC, Manassas, VA, CRL-1740) and 22Rv1 cells (ATCC, Manassas, VA, CRL-2505) were maintained in Roswell Park Memorial Institute medium (RPMI-1640, ATCC, Manassas, VA, USA) with 10% fetal bovine serum (FBS) (Thermo Fisher Scientific, Waltham, MA, USA) and HEK-293FT cells (Invitrogen, Waltham, MA, R700-07) were cultured in Dulbecco’s Modified Eagle Medium (DMEM) with 10% FBS (Thermo Fisher Scientific, Waltham, MA, USA). Cells were authenticated by STR profiling and tested negative for Mycoplasma using Universal Mycoplasma Detection Kit (ATCC #30-1012 K).

Analyses of loss or gain of function in LNCaP and 22Rv1 cells, respectively, was performed as described in (Arriaga et al., 2024; Papachristodoulou et al., 2021). For loss of function studies, we analyzed the consequences of silencing SIRT1 in LNCaP cells using two independent shRNAs based on LT3GEPIR T3G-GFP-(miR-E)-PGK-Puro-IRES-rtTA3 (Addgene #111177) (Fellmann et al., 2013). The sequences of the shRNA are provided in the Key Resources Table. For gain of function studies, we analyzed the consequences of overexpression of SIRT1 in CRISPRa-engineered 22Rv1 cells, as described in (Arriaga et al., 2024). A non-targeting sgRNA (sgNT) and two individual sgRNAs targeting *SIRT1* (sgSIRT1 #1 and #2) were generated based on the protospacer sequences identified in the hCRISPRa-v2 library in (Horlbeck et al., 2016), and subcloned into the sgRNA template pU6-sgRNA EF1Alpha-puro-T2A-BFP (Addgene #60955). Primer sequences are provided in the Key Resources Table. Procedures for the generation and infection with lentivirus were done as in (Arriaga et al., 2024; Papachristodoulou et al., 2021).

For analyses in culture, cells were seeded at 2x10^5^ cells (22RV1) or 5x10^5^ cells (LNCaP) per 10 cm dish in RPMI + 10% FBS. The following day, cells were changed to RPMI phenol red-free + 5% charcoal-stripped serum (CSS) and grown in the DMSO and ethanol or 1 mM dibutyryl cyclic-AMP (db-cAMP, STEMCELL Technologies, Vancouver, BC, Canada) and 0.5 mM 3-Isobutyl-1-methylxanthine, IBMX (Thermo Fisher Scientific, Waltham, MA, USA).

Colony formation and proliferation assays were performed as described in (Papachristodoulou et al., 2021). Briefly, for colony formation assays, 22RV1 cells (1x10^3^ cells) were seeded in triplicate in 6-well plates and grown for 12 days. Colonies were visualized by crystal violet staining and quantified using ImageJ. To measure proliferation, we performed 3-(4,5-dimethylthiazol-2-yl)-2,5-diphenyltetrazolium bromide (MTT) proliferation assays. 1x10^4^ cells were seeded in triplicate in 96-well plates and grown up to 72h. MTT-based proliferation was quantified by measuring OD at 560 nm in a Varioskan LUX multimode microplate reader (Thermo Fisher Scientific, Waltham, MA, USA).

For analyses of tumor growth in vivo, 1x10^6^ 22RV1 cells in 30 µL of 50/50 PBS/Matrigel (Corning, Corning, NY, USA) were injected into the anterior prostate (AP) of immunodeficient Athymic Nude mice (Envigo, Indianapolis, IN, USA) as described (Arriaga et al., 2024). Mice were euthanized at 40 days post-injection and tumors were collected and analyzed as described (Arriaga et al., 2024).

For pharmacological inhibition of SIRT1, we used a selective SIRT1 inhibitor, Selisistat EX-527 (Purity: 99.78% from Selleckchem) resuspended in DMSO (Adams et al., 2024; Napper et al., 2005). LNCaP cells were seeded at 5 x 10^5^ cells per 60 mm dish in RPMI + 10% FBS. The following day, cells were changed to RPMI phenol red-free + 5% charcoal-stripped serum (CSS) and treated with either DMSO and ethanol, or 1 mM db-cAMP (from 500 mM stock in 50/50 in Ethanol/PBS) (STEMCELL Technologies, Vancouver, BC, Canada), 0.5 mM 3-Isobutyl-1-methylxanthine, IBMX (from 1M stock in DMSO) (Thermo Fisher Scientific, Waltham, MA, USA), or Selisistat EX-527 (from 50 mM stock in DMSO) (Selleckchem). To determine the dose of Selisistat that inhibits SIRT1 activity in cultured LNCaP cells, we examined the overall acetylation of proteins using anti-acetyl Lysine antibodies (Abcam ab22550) (Key Resources Table).

For all cell culture studies, validation of mRNA expression levels was performed by quantitative real-time PCR with QuantiTect SYBR Green PCR Kit (QIAGEN, Hilden, Germany), using GAPDH as an internal control. Relative expression levels were calculated using the 2^-ΔΔCT^ method, as described (Papachristodoulou et al., 2021). Sequences of the primers are provided in the Key Resources Table. Validation of protein levels was performed by Western blot using total cell lysates extracted with RIPA buffer, as described (Papachristodoulou et al., 2021). Antibodies described in the Key Resources Table.

### Statistical analysis

Statistical analysis was performed using Graphpad Prism software (version 9.4.1) and R Studio (v1.3.1093), R (v4.1.2). Survival curves for the *Sleeping Beauty* mice were plotted based on Kaplan-Meier analysis and significance was calculated based on the Log-rank Test using the *survival* (3.5-5) and *survminer* (0.4.9) packages in R. Metastasis and NEPC frequencies were calculated and significance was evaluated based on the Fisher’s Exact Test in R. Tumor weight of *SB* tumors was analyzed and significance was calculated based on Welch’s two sample t-test in R. Analysis of Ki67 staining was performed using Graphpad Prism (version 9.4.1) and significance was calculated with one-way ANOVA with Tukey’s multiple comparisons test. Functional validation experiments were plotted using Graphpad Prism and significance was calculated based on one-way ANOVA with Dunnett’s multiple comparisons test for tumor weights, mRNA relative expression from cell lines, relative number of colonies from colony formation assay and OD from MTT assay. Significance for the above-mentioned statistical tests was assumed for *P* < 0.05.

### Data accession

The RNA sequencing expression profiling data have been deposited in the Gene Expression Omnibus (GEO) database (https://www.ncbi.nlm.nih.gov/geo/) with the following accession codes: **GSE271066** (a description of all data from the study), **GSE271054** (LM-PCR-based targeted genomic sequencing from NPp53-SB(+) mice), **GSE271053** Mouse RNASeq data from NP/NPp53-SB(—) and NP/NPp53-SB(+) mice.

## Additional info

### Financial support for each author

This work was supported by grants R01CA173481, R01CA283068, and R01CA183929 to CAS, R01CA251527 to MMS, P01CA265768-03 to MMS and CAS, the NCI’s Center for Cancer Systems Therapeutics (CaST) award U54CA274506 to AC and CAS, the NCI Outstanding Investigator award R35 CA197745 to AC, and the NIH Shared Instrumentation Grants S10 OD012351, S10 OD021764 and S10OD032433 to A.C., a Prostate Cancer Challenge Award to MMS and AC, and a Prostate Cancer Challenge Award to MAR. FNA was supported by a 2020 AACR-AstraZeneca Stimulating Therapeutics Advancements through Research Training (START) Grant (20-40-12-NUNE). AV was supported by a U.S. Department of Defense Early Investigator Research Award (W81XWH19-1-0337) and an Early Career Development Pilot Award NIH/NCI Cancer Center, funded through the Cancer Center Support grant, P30CA013696. MZ was supported in part by the National Center for Advancing Translational Sciences, National Institutes of Health, Grant Number UL1TR001873. JMA was supported by the Dean’s Precision Medicine Research Fellowship from the Irving Institute for Clinical and Translational Research at Columbia University Irving Medical Center (UL1TR001873), and a Prostate Cancer Foundation Young Investigator Award. ALEW was supported by NIDDK T35 grant (T35DK093430). C.A.S. is an American Cancer Society Research Professor supported in part by a generous gift from the F.M. Kirby Foundation.

## Acknowledgements

We are indebted to Timothy Starr, the primary architect of the TAPDANCE software, for his assistance and insights in applying it to our Sleeping Beauty screen. Some parts of Figures 1, 2, 4, 5 and 6 were created with <u>BioRender.com</u> using an institutional license sponsored by Columbia University’s VP&S Office for Research. These studies were supported by Flow Cytometry and Genomics and High Throughput Screening Core facilities, which are funded in part through Herbert Irving Comprehensive Cancer Center support Grant P30 CA013696.

This work was supported by grants R01CA173481, R01CA283068, and R01CA183929 to CAS, R01CA251527 to MMS, P01CA265768-03 to MMS and CAS, the NCI’s Center for Cancer Systems Therapeutics (CaST) award U54CA274506 to AC and CAS, the NCI Outstanding Investigator award R35 CA197745 to AC, and the NIH Shared Instrumentation Grants S10 OD012351, S10 OD021764 and S10OD032433 to A.C., a Prostate Cancer Challenge Award to MMS and AC, and a Prostate Cancer Challenge Award to MAR. FNA was supported by a 2020 AACR-AstraZeneca Stimulating Therapeutics Advancements through Research Training (START) Grant (20-40-12-NUNE). AV was supported by a U.S. Department of Defense Early Investigator Research Award (W81XWH19-1-0337) and an Early Career Development Pilot Award NIH/NCI Cancer Center, funded through the HICCC Cancer Center Support grant, P30CA013696. MZ was supported in part by the National Center for Advancing Translational Sciences, National Institutes of Health, Grant Number UL1TR001873. JMA was supported by the Dean’s Precision Medicine Research Fellowship from the Irving Institute for Clinical and Translational Research at Columbia University Irving Medical Center (UL1TR001873), and a Prostate Cancer Foundation Young Investigator Award. ALEW was supported by NIDDK T35 grant (T35DK093430). C.A.S. is an American Cancer Society Research Professor supported in part by a generous gift from the F.M. Kirby Foundation.

## Legend

**CIS FDR:** adjusted p-value using false discovery rate for each common-insertion site from TAPDANCE analysis

**DGE FDR:** adjusted p-value using false discovery rate for differential gene expression between NEPC vs. non-NEPC tumors

**CINDy FDR:** adjusted p-value using false discovery rate for each candidate modulator gene from CINDy analysis

**VIPER FDR:** adjusted p-value using false discovery rate for master regulators from VIPER analysis

**DGE logFC:** logFold-Change between NEPC vs. non-NEPC tumors from differential gene expression analysis

**VIPER Score:** VIPER score from differential protein activity analysis between NEPC vs. non-NEPC tumors

## Supplementary Materials

**A forward genetic screen identifies Sirtuin1 as a driver of neuroendocrine prostate cancer** Nunes de Almeida et al.

## Supplementary Figures (attached)

**Figures S1.**
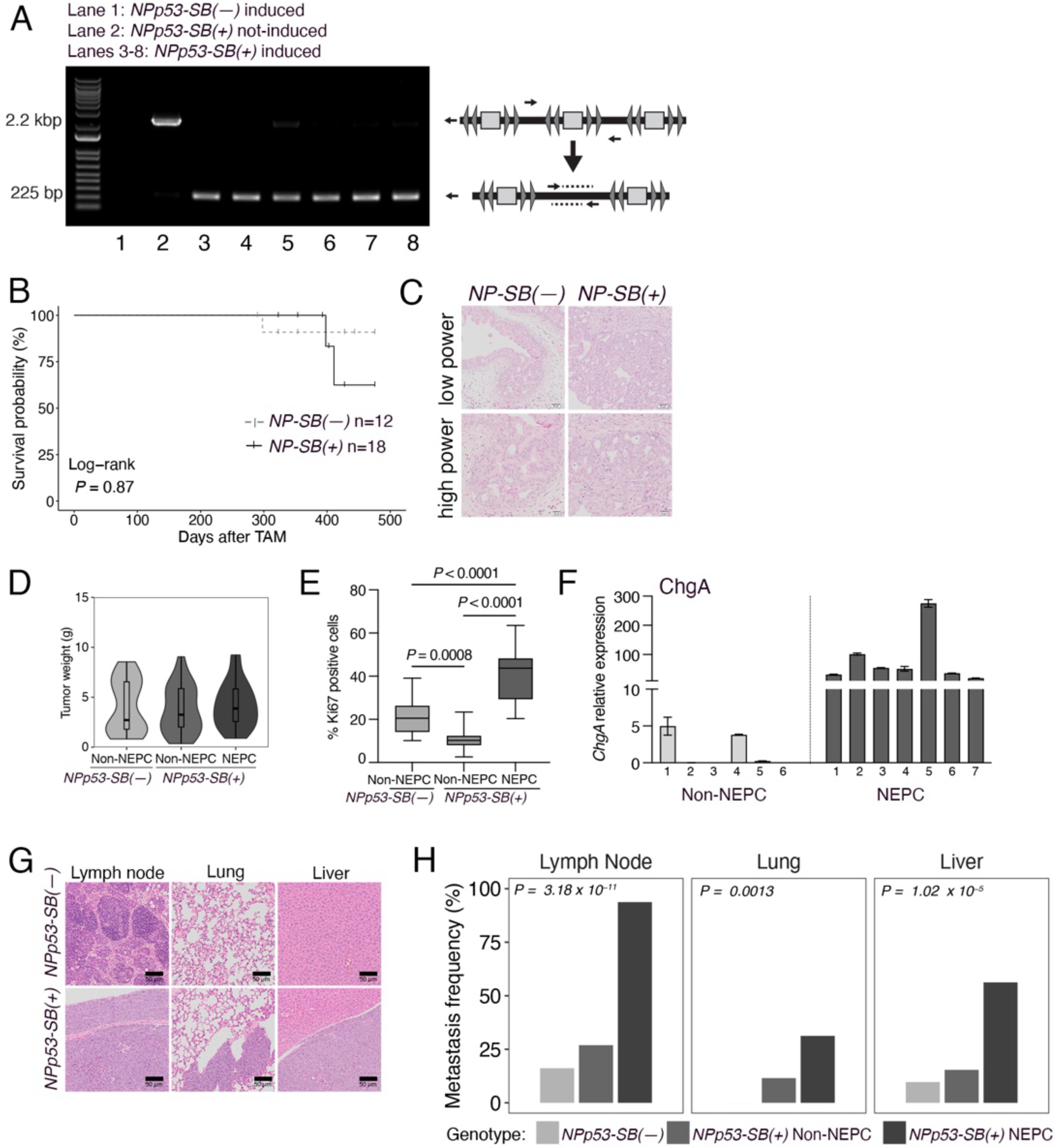
Additional analyses of the prostate phenotype of *SB* mice. A. PCR analyses showing excision of the *T2/Onc2* transposon in *NPp53-SB(+)* induced tumors. **B,C**. Analysis of the phenotype of *NP-SB(–)* and *NP-SB(+)* mice. **B.** Kaplan Meier survival analysis. *P* value was calculated using a Log-rank test. **C.** Representative H&E images of prostate tumors showing low power and high-power. Scale bars represent, 50 µm for low power and 20 µm for high power H&E. **D-F**. Additional analyses of non-NEPC and NEPC tumors from *NPp53-SB(–)* and *NPp53-SB(+)* mice. **D.** Summary of tumor weights from *NPp53-SB(–)* (n=29) and *NPp53-SB(+)* non-NEPC (n=43) and NEPC (n=27). **E**. Quantification of Ki67 expression in *NPp53-SB(–)* and *NPp53-SB(+)* non-NEPC and NEPC (counted a minimum of 8,000 cells from 5 sections per tumor from 3 independent tumors per condition). **F**. Quantitative real-time PCR showing expression levels of chromogranin A in non-NEPC (n=6) and NEPC (n=7) *NPp53 (SB+)* tumors. **G,H**. Analyses of metastasis in *NPp53-SB(–)* and *NPp53-SB(+)* mice. **G**. Representative H&E images. Scale bars represent, 50 µm for H&E. **H**. Summary of metastasis frequency in the indicated tissues. Detailed analyses of the metastasis phenotype is provided in Table S1.

**Figures S2.**
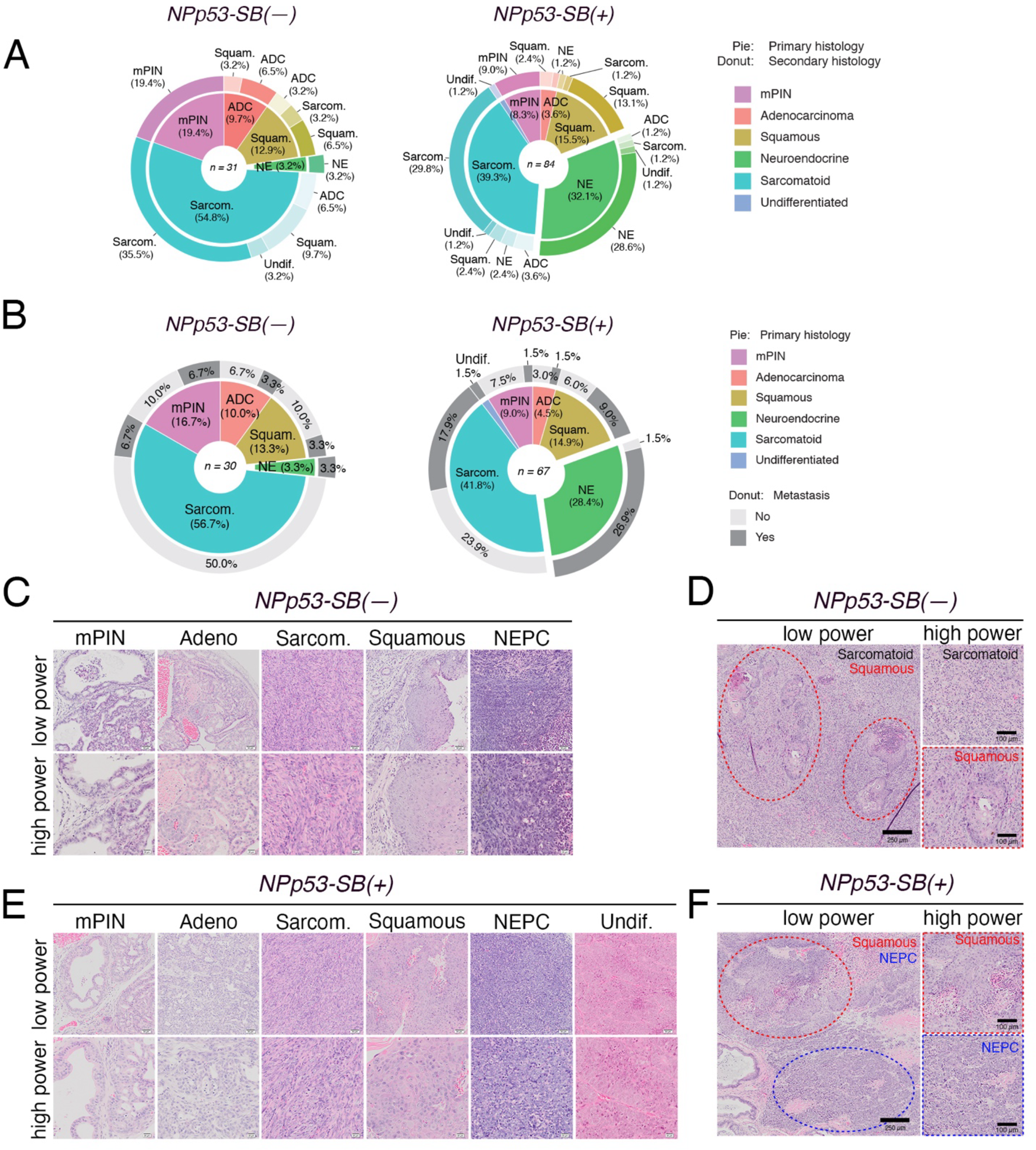
Detailed analyses of the histopathological and metastasis phenotype in the *NPp53-SB(–)* and *NPp53-SB(+)* mice. **A,B.** Distribution of histology phenotypes and metastasis incidence in *NPp53-SB(–)* (n=30) and *NPp53-SB(+)* (n=67) mice. Histology phenotypes were classified as mouse prostatic intraepithelial neoplasia (mPIN), adenocarcinoma (ADC), squamous (Squam), neuroendocrine (NE), sarcomatoid (Sarcom) or undifferentiated (Undif) **A.** Distribution of primary histology (pie chart) and secondary histology (donut chart). **B.** Distribution of primary histology (pie chart) and metastatic status (donut chart). **C-F**. Representative H&E images showing each of the observed tumor phenotypes and their heterogeneity. **C-D.** Selected *NPp53-SB(–)* cases. **E-F.** Selected *NPp53-SB(+)* cases. Scale bars represent, 50 µm for low power (top) and 20 µm for high power (bottom) images (**C** and **E**), 250 µm for low power (left side) and 100 µm for high power (right side) (**D** and **F**). Detailed histopathology analysis is provided in Table S1.

**Figures S3.**
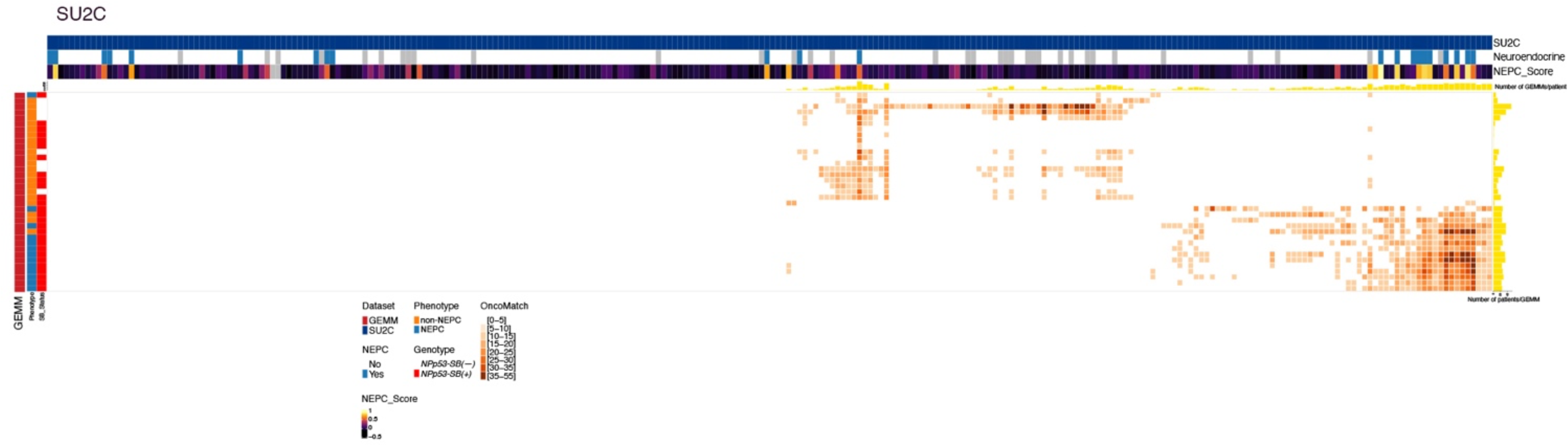
Additional OncoMatch analyses. OncoMatch heatmap representing conservation of master regulators (MRs) between individual *SB* tumors (rows, n=32) and patient tumors (columns, n=266) from the SU2C cohort. Patient variables (*i.e.* assessment of neuroendocrine features and NEPC features) are shown on the top three bars, while GEMM variables (*i.e.* phenotype and genotype) are shown on the left side vertical bars. Yellow bars at the top of the heatmaps show the number of GEMMs that match each patient, while the yellow bars on the right side show the number of patients that match each GEMM.

**Figures S4.**
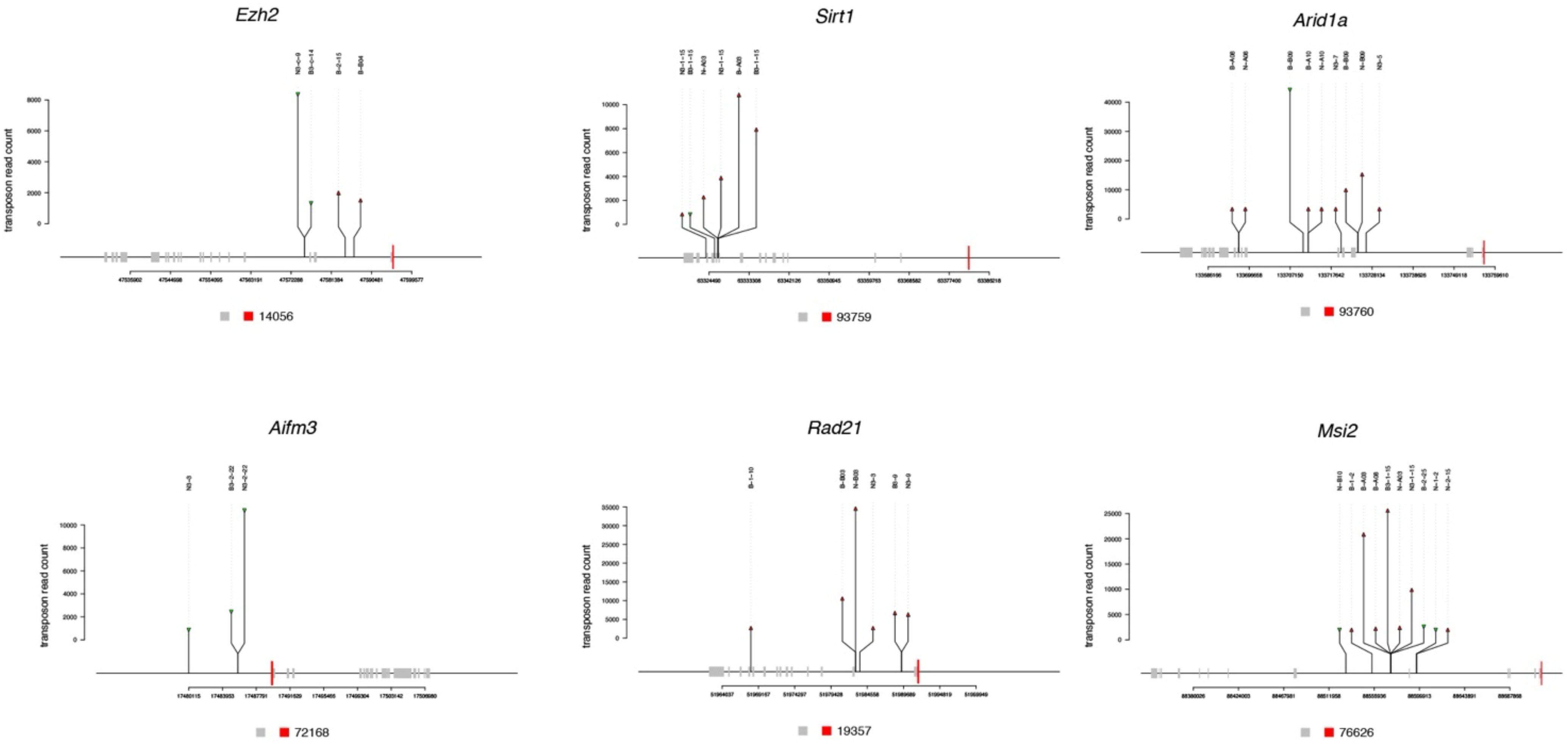
Lollipop representation of selected CIS-associated genes. Shown are examples of the genomic structure of selected CIS-genes (Ezh2, Sirt1, Arid1a, Aifm3, Rad21, and Msi2) with promoters represented in red and exons in grey. Vertical arrows indicate individual transposon insertion sites, with the arrow height proportional to the transposon read count at that location. Red arrows represent insertions in the same transcriptional direction as the gene, suggesting potential transcriptional activation or alternative isoform expression. Green arrows indicate insertions in the opposite direction, potentially disrupting gene function.

**Figures S5.**
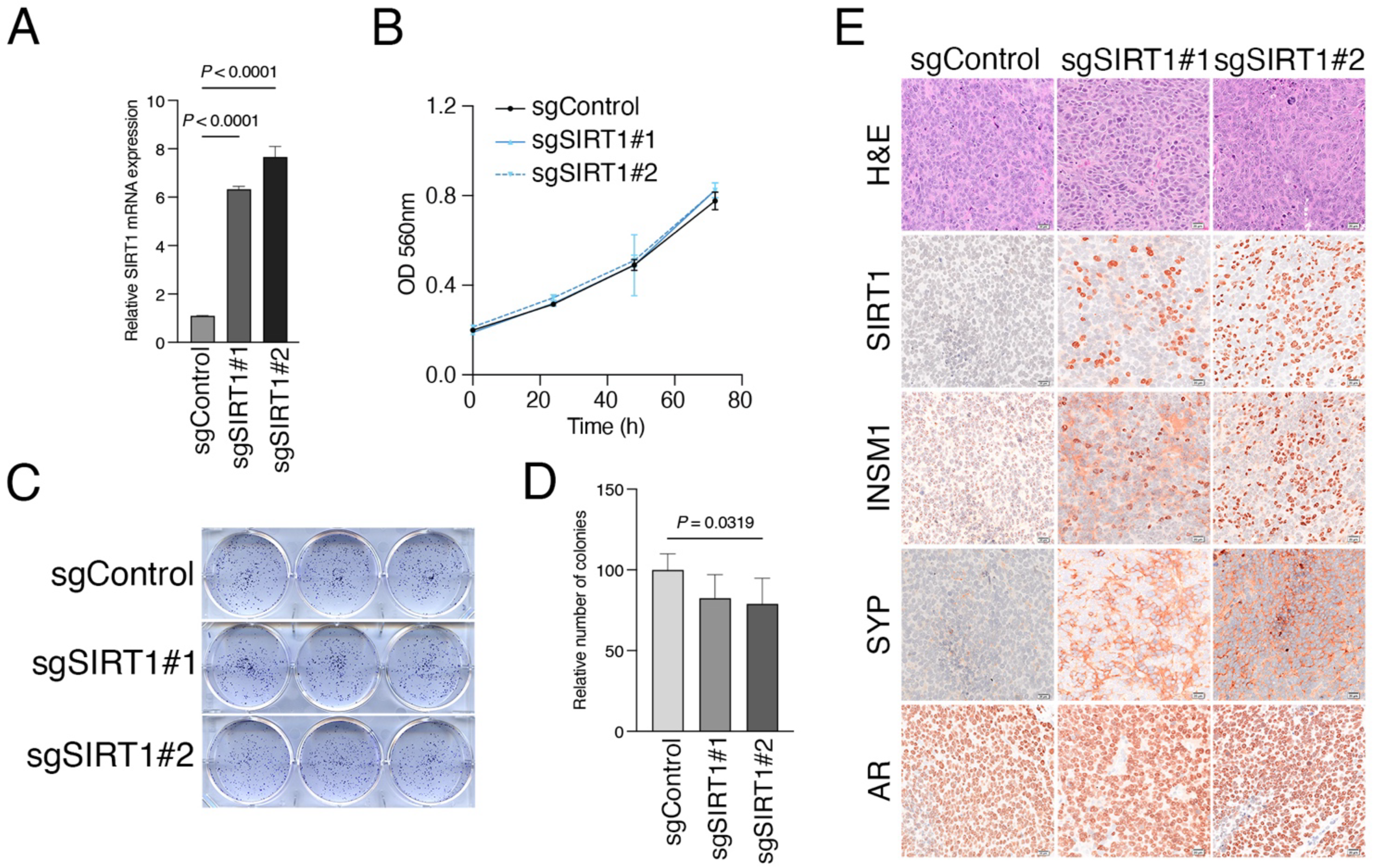
Additional analyses of validation studies. **A**. Real-time PCR showing relative expression of SIRT1 in 22RV1 cells. **B**. Quantification of cell proliferation by detection of MTT absorbance. OD, optical density. **C,D**. Crystal violet staining of colony formation assay showing 22RV1 CRISPRa cell lines treated for 12 days with sgControl, sgSIRT1 #1 or #2, and quantification of the number of colonies. **E.** Representative images of the 22RV1 prostate tumors. Shown are hematoxylin and eosin (H&E) and immunostaining for staining for SIRT1, INSM1, Synaptophysin (SYP), and AR. Scale bars represent 20 µm.

Key Resources Table (uploaded separately)

## Supplementary Tables (uploaded separately)

Table S1: Summary of *Sleeping Beauty* mouse phenotypes. Detailed description of individual mice analyzed for RNA sequencing, identification of common insertion sites (CIS) and histopathology

Table S2: Differential gene expression analyses. Comparison of RNA seq profiles from *SB* mice with NEPC or non-NEPC

Table S3: Protein activity signatures from mouse *SB* tumors.

Table S4: Protein activity of cluster analyses of mouse *SB* tumors

Table S5: OncoMatch analyses comparing *SB* tumors and human patient cohorts

Table S5A: OncoMatch comparing *SB* tumors and SU2C

Table S5B: OncoMatch comparing *SB* tumors and Beltran Table S5C: SU2C Protein Activity

Table S5D: Beltran Protein Activity

Table S6: SB-barcodes Barcodes used to sequence each *SB* tumor sample.

Table S7: Summary of CIS analyses and CIS-associated genes

Table S7A: List of common insertion sites (CIS) determined by TAPDANCE analysis, occurrence in *SB* tumors with NEPC and prediction for driving of transcription

Table S7B: List of CIS-associated genes determined by TAPDANCE analysis, occurrence in SB tumors with NEPC and prediction for driving of transcription

Table S8: Prioritization of CIS-associated genes. Integration of TAPDANCE, differential gene expression (DGE) and CINDy analyses to prioritize CIS-associated genes and identify candidate drivers of NEPC.

